# Pollen sequencing reveals barriers and aberrant patterns of recombination in interspecific tomato hybrids

**DOI:** 10.1101/2022.07.02.498571

**Authors:** Roven Rommel Fuentes, Ronald Nieuwenhuis, Jihed Chouaref, Thamara Hesselink, Willem van Dooijeweert, Hetty C. van den Broeck, Elio Schijlen, Paul Fransz, Maike Stam, Hans de Jong, Sara Diaz Trivino, Dick de Ridder, Aalt D.J. van Dijk, Sander A. Peters

## Abstract

Tomato is the most consumed vegetable in the world. Increasing its natural resistance and resilience is key for ensuring food security within a changing climate. Plant breeders improve those traits by generating crosses of cultivated tomatoes with their wild relatives. Specific allele introgression relying on meiotic recombination, is hampered by structural divergence between parental genomes. However, previous studies of interspecific tomato hybridization focused in single cross or lacked resolution due to prohibitive sequencing costs of large segregating populations. Here, we used pooled-pollen sequencing to reveal unprecedented details of recombination patterns in five interspecific tomato hybrids. We detected hybrid-specific recombination coldspots that underscore the influence of structural divergence in shaping recombination landscape. Crossover regions and coldspots show strong association with specific TE superfamilies exhibiting differentially accessible chromatin between somatic and meiotic cells. We also found gene complexes associated with metabolic processes, stress resistance and domestication syndrome traits, revealing undesired consequences of recombination suppression to phenotypes. Finally, we demonstrate that by using resequencing data of wild and domesticated tomato populations, we can screen for alternative parental genomes to overcome recombination barriers. Overall, our results will allow breeders better informed decisions on generating disease-resistant and climate-resilient tomato.

## Introduction

Crop breeding relies on the availability of genetic diversity to generate novel allele combinations that are agronomically valuable. However, long term selection by inbreeding often causes loss of essential allelic information. To reintroduce lost genetic variation, breeders have introgressed alien chromatin by crossing crops with wild relatives followed by repeated backcrossing and selection. Among the most desirable traits to be incorporated into the breeding material are abiotic stress and disease resistance, higher yield, and fruit quality ^1^. The success of introgression breeding largely depends on the process of recombination to introduce genetic material from the donor to the recipient crop. Meiotic recombination generates genetic diversity, but may also break apart co-adapted allele combinations, resulting in fitness reduction ^2^. Moreover, low frequency or complete absence of recombinantion in a genomic region leads to linkage drag and limits the ability of breeders to develop novel allele combinations. Chromosome regions where recombination is suppressed are found in pericentromeres, including retrotransposons and other DNA-methylated regions ^3,4^. Furthermore, heterozygous structural variants (SVs) have been reported to limit pairing and crossovers (COs) or lead to lethal gametes, suggesting that genomic rearrangements affect recombination patterns, especially in hybrids ^5–8^.

Genomic rearrangements may exist between related species and different genotypes of the same species. Characterization of these rearrangements have revealed recombination coldspots, some of which are associated with resistance genes or adaptive traits ^9,10^. Due to absent or diminished COs in SV regions, clusters of tightly linked alleles known as supergenes are inherited together, contributing to local adaptation and reproductive isolation ^11–13^. Suppression or absence of recombination has been found essential in speciation and domestication by allowing the fixation of alleles such as those within selective sweeps ^14,15^. One of the best studied rearrangements in plants is the 1.17Mb paracentric inversion in Arabidopsis, which shows complete lack of recombination in the rearranged genomic segment linked with fecundity under drought ^16^. It was reported that recombination is prevented by SVs in genomic regions causing self-incompatibility in Brassicaceae plants ^17^ and reproductive isolation in monkeyflower ^18^. In the backcross descendants of a *Solanum habrochaites* introgression into cultivated tomato (*S. lycopersicum*), an inversion containing the *Ty-2* resistance genes and at least 35 more genes causes linkage drag, unabling selection of appropriate agronomic trait combinations in the offspring ^19,20^. Another example is the lack of CO in the inverted region of a *S. esculentum* x *S. peruvianum*, containing the nematode-resistance gene, Mi-1, and other genes conferring resistance to different pathogens and insects ^5^.

Although previous studies addressed the role of SVs as recombination barriers in a limited number of genomic regions, a genome-wide analysis of decreased or absent COs related to SVs in tomato and multiple hybrid crosses is currently lacking, due to the absence of cost-effective and high-resolution crossover detection methods and accurate SV prediction. The effect of structural differences on chromosome pairing during meiosis has been studied using electron microscopy ^21^, comparing spreads of synaptonemal complexes (SCs) from multiple F1 tomato hybrids. Moreover, the synaptic configurations revealed mismatched kinetochores, inversion loops and translocation complexes, pointing to structural differences in the parental genomes that likely influence the recombination landscape. However, their electron microscopic studies could not reveal the consequences of erratic recombination patterns between the parental partners. To analyze these patterns in higher resolution, Demirci, et al. ^22^ sequenced F6 recombinant inbred lines obtained from a cross between tomato (*S. lycopersicum*) and its wild relative *S. pimpinellifolium*, and computationally detected recombination sites. We subsequently developed a less laborious and costly method, involving pollen profiling. Using a pool of pollen from *S. pimpinellifolium* x *S. lycopersicum* F1 hybrids, we generated a recombination landscape at nucleotide resolution level ^23^, revealing significant reduction of COs in heterozygous deletions ^15^.

To better understand occurrence and frequencies of CO events, we profiled here the recombination landscape in multiple crosses of tomato and wild relatives by sequencing pools of pollen gametes. We identified CO coldspots in each hybrid cross and examined recombination patterns and barriers. Our results suggest a major role for SVs and transposable elements in shaping the recombination landscape in hybrids, specifically in suppressing COs in gene complexes that relate to adaptation, speciation, and domestication. In addition, we present an example of syntenic and non-syntenic accessions for specific genomic regions, which may be considered in the selection of parental breeding lines as so called ‘bridge accessions’ to avoid or overcome introgression bottlenecks.

## Results

### Crossovers in multiple hybrid crosses

In this study, we have generated hybrid crosses of *S. lycopersicum Heinz1706* and its wild relatives *S. pimpinellifolium* (CGN14498; **PM**)*, S. neorickii* (LA0735; **NE**), *S. chmielewskii* (LA2663; **CH**)*, S. habrochaites* (LYC4; **HB**), and *S. pennelli* (LA0716; **PN**). Hereafter, we use these abbreviations and the species name when referring to the hybrids and the parental genome, respectively. The pool of pollen from each hybrid was sequenced using 10X Genomics kits (**Supplementary Table 1**) based on the protocol described in Fuentes, et al. ^23^. To detect crossover events (COs), we first profiled single-nucleotide polymorphism (SNP) markers. We then filtered out regions prone to false positive COs, manifested by a high density of heterozygous SNPs and excessive sequence coverage (**Supplementary Figure 1**). *S. pennellii* and *S. pimpinellifolium* were the most distant and closest species to *S. lycopersicum* in this study and has the highest and lowest number of SNPs with respect to the reference genome (*S. lycopersicum;* SL4.0), respectively (**Table 1**). Using the filtered SNPs, we were able to detect haplotype shifts, leading to identification of putative recombinant haplotypes. These were further screened as described in the Methods (**Supplementary Figure 2**).

**Table 1.**
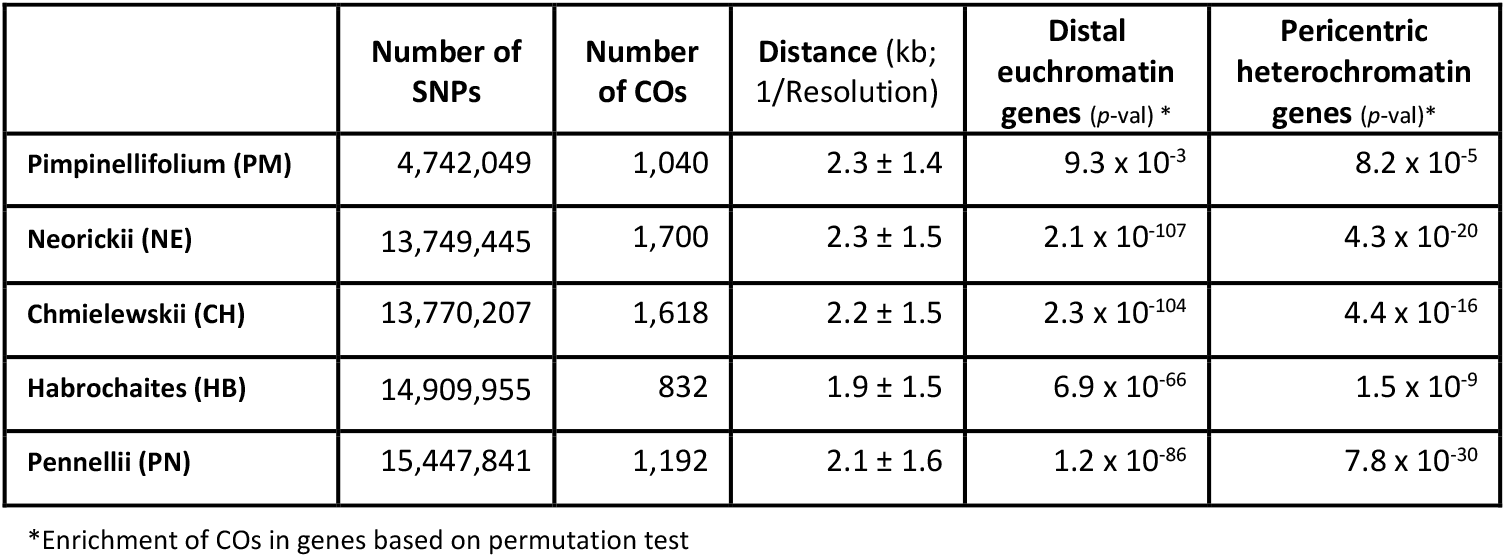
Crossovers detected in multiple interspecific (with *S. lycopersicum*) hybrid populations

|  | Number of SNPs | Number of COs | Distance (kb; 1/Resolution) | Distal euchromatin genes (*p*-val) * | Pericentric heterochromatin genes (*p*-val) * |
| --- | --- | --- | --- | --- | --- |
| Pimpinellifolium (PM) | 4,742,049 | 1,040 | 2.3 ± 1.4 | 9.3 × 10<sup>-3</sup> | 8.2 × 10<sup>-5</sup> |
| Neorickii (NE) | 13,749,445 | 1,700 | 2.3 ± 1.5 | 2.1 × 10<sup>-107</sup> | 4.3 × 10<sup>-20</sup> |
| Chmielewskii (CH) | 13,770,207 | 1,618 | 2.2 ± 1.5 | 2.3 × 10<sup>-104</sup> | 4.4 × 10<sup>-16</sup> |
| Habrochaites (HB) | 14,909,955 | 832 | 1.9 ± 1.5 | 6.9 × 10<sup>-66</sup> | 1.5 × 10<sup>-9</sup> |
| Pennellii (PN) | 15,447,841 | 1,192 | 2.1 ± 1.6 | 1.2 × 10<sup>-86</sup> | 7.8 × 10<sup>-30</sup> |

\*Enrichment of COs in genes based on permutation test

We detected a total of 6,382 COs in all hybrids, mostly located in distal segments of chromosomes, which is consistent with previous reports in tomato and other plant species ^3,8^. For each hybrid, COs are confined to 6-12% and below 1% of the distal euchromatin (DEU) and pericentric heterochromatin (PCH), respectively, and in total CO regions account for only 2% of the whole genome, consistent with other eukaryotic organisms where recombinations are concentrated in hotspots ^3,8,24,25^. Although PCH regions of tomato are known to exhibit low recombination rates ^26,27^, we detected there a total of 710 COs (11.1%) from all hybrids, which most likely locate in euchromatin island in PCH. It was proposed that suppression of double-strand-breaks (DSBs), the precursor of COs, by heterochromatin on repetitive DNA helps safeguard against genome destabilization ^27,28^.

We validated the resulting recombination profile of PM by comparison to existing CO data. The COs in the pollen gametes significantly overlap with COs previously detected in a RIL population of the same parental cross (Fisher’s exact test; P = 5.8 x 10^-18^) ^22^. More in detail, the frequency of COs in sliding genomic windows in DEU or in PCH also revealed a significant correlation between the recombination landscapes generated from pollen and the RIL population sequence data (Spearman’s rank correlation; distal euchromatin, ρ = 0.33; P < 2.2 x 10^-16^; pericentric heterochromatin, ρ = 0.22; P < 2.2 x 10^-16^). Furthermore, comparison with historical recombination hotspots detected in natural populations of wild and domesticated tomato ^15^ revealed that the COs in hybrids overlap with 294 (Fisher’s exact test; P = 2.0 x 10^-13^) and 36 (Fisher’s exact test; P = 3.8 x 10^-11^) historical hotspots in DEU and PCH, respectively. Previous observations and our results both confirm recombination sites in PCH, which thus far were rarely observed due to their low frequency and the limitations of other CO-detection methods.

The vast majority (5,150) of COs are located within genes and their 1kb flanking regions, while another 471 are positioned between 1kb and 3kb from genes (**Table 1**; **Figure 1a**; **Supplementary Figure 3**). PM and NE have the lowest CO resolution, defined as the inverse of the distance between the SNP markers bounding the CO region (resolution = 1/distance). We consider a detected CO as high resolution if the distance between the markers flanking the CO region is below 1kb, *i.e*. if the resolution is above 0.001. The number of COs near or within genes, in both DEU and PCH regions, is significantly higher than expected by chance (**Table 1)**. Although genic regions account for only 15.5% of tomato genome, majority of the CO regions overlap gene features (**Figure 1a**). *S. pimpinellifolium* COs apparently overlap more with intergenic regions than COs in other hybrids. One possible confounding factor could be that *S. pimpinellifolium* has fewer SNPs than the other species (**Table 1**), which may lead to a lower resolution of detected COs. To determine if this contributed to the higher overlap between gene features and COs in PM, COs with similarly distributed resolution for all hybrids were separately analyzed (**Supplementary Figure 4A**). However, the result still shows the same higher intergenic overlap of crossover events in PM (**Supplementary Figure 4B**), which is unexpected and enigmatic.

**Figure 1.**
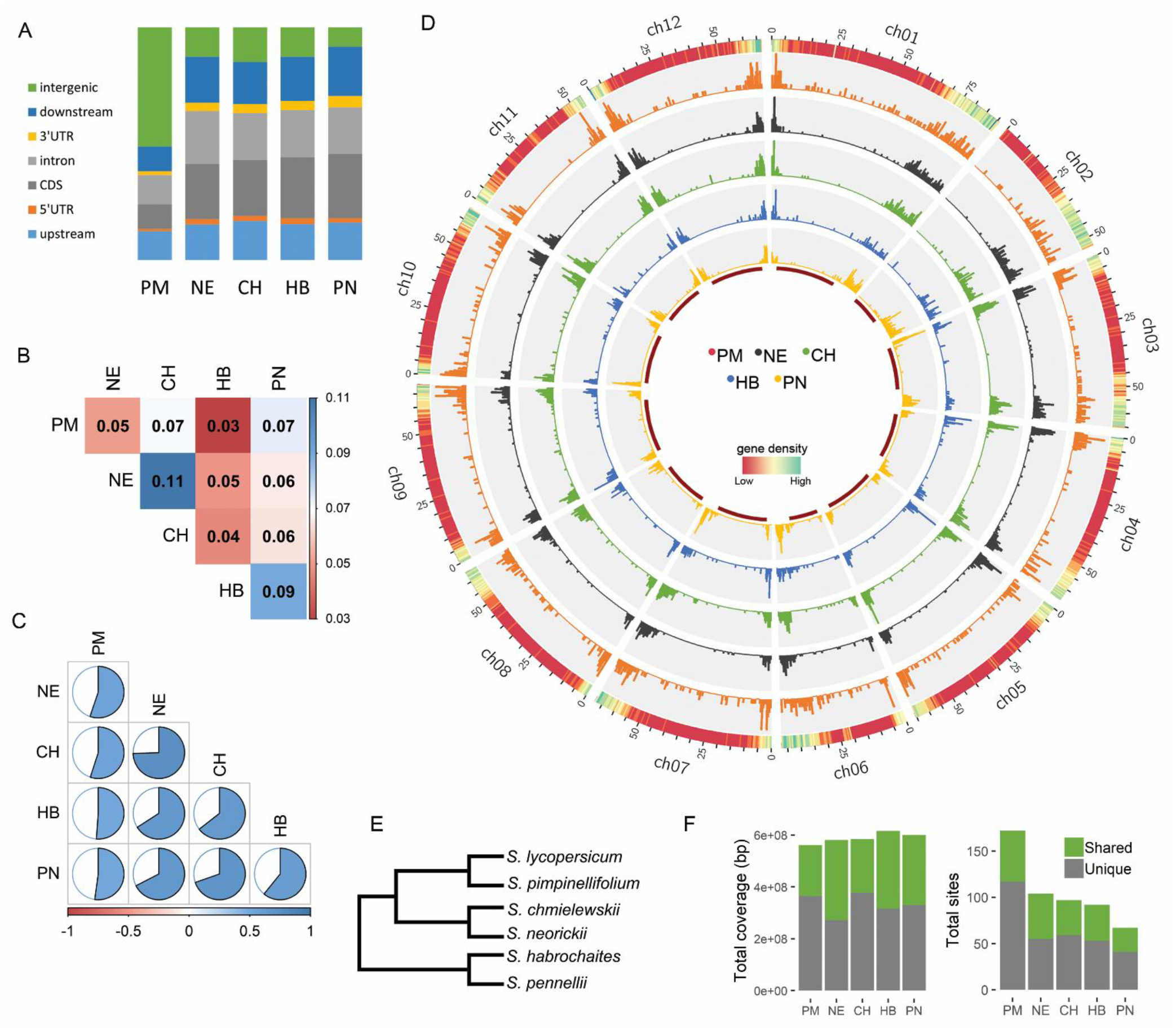
Recombination landscape. A) Distribution of crossover regions over gene features. Upstream and downstream covers 1 kb from the transcription start and termination sites, respectively. B) Fraction of shared CO sites and C) Correlation of the genome-wide CO landscape between hybrids. D) Distribution of COs per hybrid. The outermost track indicates gene density while the red innermost track marks the pericentric heterochromatin regions. E) Phylogenetic tree of the parental species based on Moyle ^96^ F) Coverage and number of recombination coldspots in different crosses.

Aside from association with genes, sequence motifs are discovered in CO regions too ^22,29,30^. Using high resolution COs, we found that CTT-like repeats, poly-AT and A-rich motifs are actually enriched in regions flanking rather than within COs (**Supplementary Figure 5**). These CO motifs have been previously identified in other plant species but due to low resolution, it was not possible to determine whether they were actually located within or just near CO regions. However, their close proximity to crossover sites hints that they may have a role in recruiting recombination-promoting factors as previously proposed ^31^.

### Unique recombination patterns between hybrids

All hybrids show similar recombination landscapes with COs mostly at distal, gene-rich chromosome regions. Yet, there are also unique, local patterns of COs shown in **Figure 1d**. Comparisons of recombination profiles from different hybrids are essential to learn about variability and genomic factors contributing to CO patterns. To examine similarities between hybrids, we first identified overlapping COs between hybrids and found a significantly higher fraction than expected by chance (**Figure 1b**). Among all pairs, the highest overlap of COs is observed between hybrids with wild parents that are evolutionarily closely related to each other (NE and CH or HB and PN). On the contrary, CO sites in PM have more overlap with PN than with other closely related species, which does not reflect their evolutionary distance (**Figure 1e**). A low but significant overlap has also been observed when comparing recombination hotspots in natural populations of wild and domesticated rice, cocoa and tomato ^15,32,33^. About 35.3% (1,161) of CO regions, containing a total of 3,996 crossover events, are shared by at least two hybrids. CO regions per hybrid cover around 2% of the genome, whereas they cover 10% (77.6 Mbp) when combined, apparently not extensively overlapping, thus indicating divergent CO regions between the hybrids.

Given the low rate of CO region overlap between hybrids, we decided to investigate whether the overall recombination landscape across the genome is significantly correlated between hybrids. **Figure 1c** shows that NE and CH have the most similar landscape. The low CO overlap (4%; **Figure 1b**) between CH and HB does not translate to a low landscape correlation (r^2^ = 0.64); similarly, despite the high overlap between PM and PN COs (7%), the correlation coefficient of their landscapes is one of the lowest among all pairs (r^2^ = 0.52), consistent with their evolutionary distance. Although the number of overlapping COs is significant, it is far less than the non-overlapping COs that contribute more to shaping the overall recombination landscape. This result suggests that despite the similar overall landscape, the hybrids exhibit local differences in CO patterns.

As shown in the landscapes, the patterns of genomic regions without recombination in the hybrids differ. To analyze these patterns, we identified CO coldspots of more than 1Mb and found that they cover 72-79% of the genomes, with the highest coverage in HB and PN. Grouping by genomic position and size, we assigned coldspots into 325 *unique* and 101 *shared* clusters (**Figure 1f**), with 63.6% of the genome (6.4Mb euchromatic; 485Mb heterochromatic) lacking CO in all five hybrids, which we refer to here as *conserved* coldspots. PM has significantly shorter coldspots than the other hybrids (pairwise Wilcoxon rank-sum test; P < 1.4 x 10^-2^) and a large number of unique coldspot regions. This divergent patterns of CO region and coldspot, confirms that hybridization of tomato with different wild parents results to variable recombination bottlenecks, uncovering the additional complexity in breeding.

### Absence of COs in structural variant heterozygosity

With the results above indicating clear variation in the occurrence of COs in the different hybrids, we speculated that large genomic rearrangements between species may underlie the varying patterns of recombination. To investigate this, we detected SVs between the parental species *S. lycopersicum* and the wild relatives. Furthermore, given that heterozygous SVs may exist in the wild species genomes, allowing the F1 hybrid to inherit an allele that is similar to the reference genome, we also genotyped SVs in the F1 hybrid pollen sequences and retained only the heterozygous ones (**Figure 2a**). Combining all parental wild species genomes, we detected 59,265 SVs with size above 50bp. We found more deletions than inversions, which may be due to either the inherently low frequency of large inversions ^34^ or the difficulty of detecting inversions compared to deletions (**Figure 2b**). Among the wild genomes, HB and PN have the highest number of SVs, which are also significantly longer compared to the other parental genomes (**Supplementary Figure 6**). To check the accuracy of the filtered SV set, we manually verified SVs from *S. pennellii* using dot plots between *S. lycopercisum* and *S. pennellii* assemblies (**Supplementary Figure 7**). 88% of 50 randomly selected deletions are supported, while an additional 10% belong to more complex translocation events and the remaining 2% are false positives. For inversions, we found 76.7% true positives.

**Figure 2.**
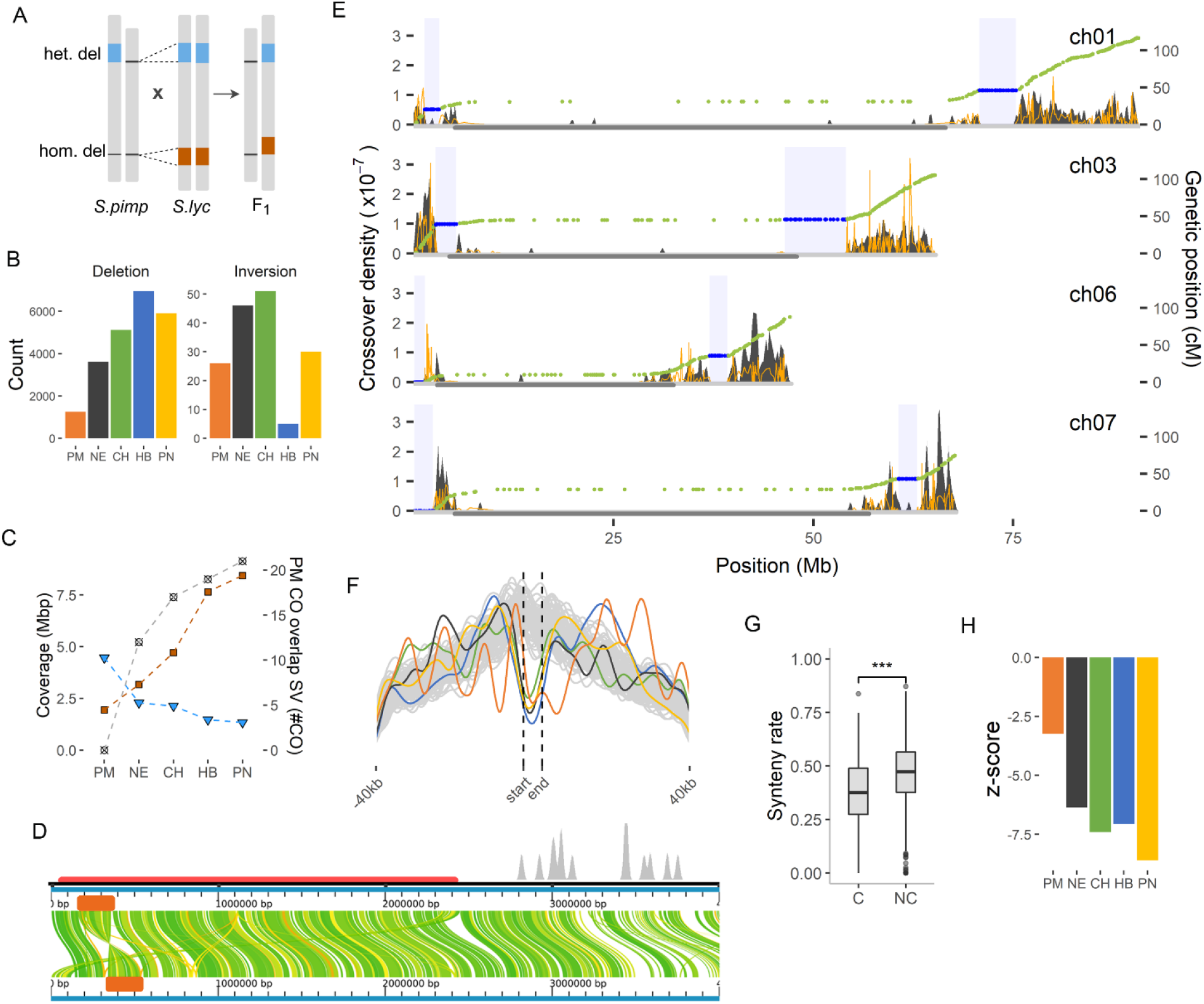
Lack of crossover in structural variations. A) Selection of parental SVs causing heterozygosity in the F1 pollen genomes. B) Frequency of SVs per wild relative. Inversions only include events > 30 kb. C) Genome coverage of COs (blue) and SVs (orange) in the PCH (left y-axis). The gray squares show the number of PM COs that overlap with SV regions in the wild genome (right y-axis). D) Large inversion (orange block) and rearrangements within the coldspot (horizonal purple segment) of chromosome 7, short arm. CO density is indicated in grey at the top. E) Crossover density of selected *S. pennellii* chromosomes (gray peaks) plotted together with Marey map (green dots) of EXPEN2012 (Sin *et al*., 2012). The blue dots are genetic markers within coldspot regions (blue box). The yellow distribution line indicates the recombination rate obtained by taking the derivative of the Marey map. The gray horizontal segment in the middle of the chromosome marks the PCH. F) Distance of COs to the nearest SV compared to the 10,000 permutation sets represented by gray lines. The vertical lines marks the boundaries of COs. G) Rate of synteny in coldspot (C) and non-coldspot (NC) regions of *S. pennellii* (Wilcoxon rank-sum test; P < 2×10^-16^). H) Suppression of COs in SV regions based on permutation test. The negative z-score means the overlap of COs in SV regions is lower than expected by chance.

To further examine the relationship between SVs and recombination landscapes, we identified rearrangements and syntenic regions between *S. lycopersicum* and *S. pennellii* assemblies and compared them against PN COs. We found that 94% of PN COs are in syntenic segments at distal chromosomal regions (Fisher’s exact test; P <0.001; **Supplementary Figure 8**), which correspond to the essential role of synteny in synapsis and crossing-over of homeologous chromosomes during meiosis ^10,35^. Using a permutation test, we indeed found strong reduction of recombination in SVs across all hybrids, specifically for SVs larger than 1kb (**Figure 2h**). Further analyses will only use SV larger than 1kb (**Supplementary Figure 9**). About 62-74% of SVs in the wild genomes overlap with coldspots, which may relate with the absence of recombination. Most SVs are located a few to tens of kilobases away from COs (**Figure 2f**), similar to *A. thaliana* ^6^, but SV size is not correlated to its distance from the CO site (**Supplementary Figure 10**).

Given that DEU and PCH in tomato have distinct genomic features, we examined their SV composition and found more SVs in DEU than in PCH regions, with an average ratio of 1.55 to 1. This agrees with the previous observations that wild and domesticated tomato accessions have higher SV density in DEU than in PCH ^34,36^. In addition, SVs in PCH are on average longer than those in DEU (Wilcoxon rank-sum test; P < 5.8 x 10^-16^; **Supplementary Figure 11**). We also observed that in PCH, higher genome coverage by SV regions comes with lower coverage by CO regions (**Figure 2c**). PM has the largest total number of CO regions in PCH, while PN has the largest number. As these PM COs overlap with the SVs in the other wild genomes, we argued that the higher SV content in other wild genomes leaves less sites for recombination in hybrids. Given that there are other complex rearrangements and SV types that we cannot detect with our data, it is likely that more divergent CO sites are defined by the presence or absence of SVs.

We identified large parts of the DEU in PN with prominent spots without CO. To validate whether these represent real coldspots, we compared them against the recombination coldspots in the EXPEN2012 linkage map ^37^. First, we identified large DEU coldspot regions in the linkage map by mapping the EXPEN2012 markers against the tomato reference genome, retrieving the physical position and subsequently plotting against the genetic position (**Figure 2e**). Then, we compared the EXPEN2012 coldspot against the PN coldspots. Large coldspots are observed in some chromosomes, spanning 0.14 to 7.64 Mb, and they match the coldspots we found in PN, demonstrating the accuracy of our method. Unlike the course-grained genetic map, the fine-scale recombination profile we generated allows comparison with genome features, aiding the elucidation of factors influencing recombination landscape.

Further inspection of these large PN coldspots revealed that they have significantly lower levels of synteny compared to non-coldspots (**Figure 2g**). These coldspots, however, may be specific to PN or may not fully overlap coldspots in other hybrids, as we have found 518 COs in the other hybrids. Among the PN coldspots, we found that at least two, specifically in the short arm of chromosomes 6 and 7, contain large inversions relative to the reference genome as previously validated using BAC-FISH ^9^. They may also correspond to the inversion loops found at the distal chromosome ends and in the euchromatin-heterochromatin borders ^21^. We were able to identify the exact location of an inversion in chromosome 7 (**Figure 2d**) by comparing genome assemblies and inspecting linked reads (**Supplementary Figure 12**). Aside from the inversion, this 2.4 Mbp coldspot region also contains other rearrangements like translocations that could inhibit proper synapsis and recombination. Upon examining the other large coldspots, we similarly found complex rearrangements and large insertions and deletions. Across all hybrids, our results suggest that SVs contributed significantly to shaping the recombination patterns by inhibiting COs, which may have been vital in the fixation of specific alleles during domestication ^15,38^. Importantly, the SVs that have been implicated with domestication of tomato can cause heterozygosity during hybridization with wild relatives which consequently suppress recombination.

### Widespread coldspots in TE regions

Aside from SVs, studies on other species also linked the presence of transposable elements (TEs) with CO incidence, specifically retrotransposons with COs suppression ^4^. In tomato hybrids, most retrotransposons (Class I), except SINEs and RTE-BovBs, indeed show suppression of COs (**Figure 3a**). However, *Stowaway* and *Tip100* (Class II), simple repeats and low complexity regions are enriched with COs. TEs associated with CO suppression are densely distributed in the PCH, whereas *Stowaway* and *Tip100* are located mostly in the DEU (**Figure 3c**). Similarly, this association with TE superfamilies was reported in historical recombination hotspots of wild and domesticated populations of tomato ^15^. As shown in **Figure 3b**, the presence of retrotransposons such as *Gypsy*, *Copia* and *L1* in a genomic region correlates with crossover suppression, consistent with the reports in many other species ^31,32,39–42^. In contrast *Stowaway* and *Tip100* show positive correlation with crossover incidence (**Supplementary Figure 13**).

**Figure 3.**
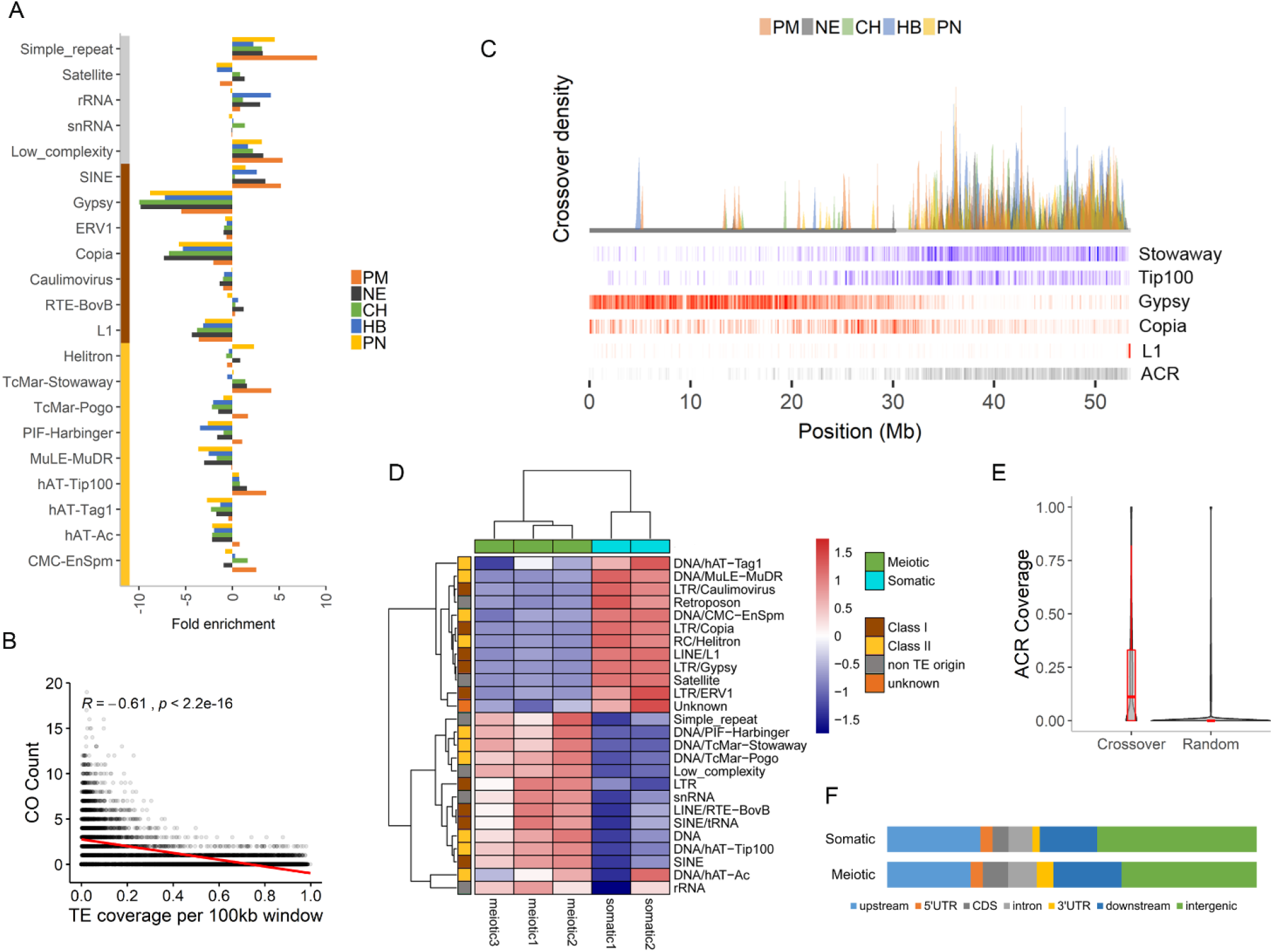
TE-associated crossovers. A) TE superfamilies and repeats showing enrichment of crossovers. Elements are clustered into DNA transposons (yellow), retrotransposons (brown) and other repeats (gray). B) Spearman’s rank correlation of crossover count and retrotransposons (Gypsy, Copia, L1) coverage in a sliding genome window. Each dot indicates a window. The red line is the local regression fitting. C) Recombination landscape of acrocentric chromosome 2 from multiple hybrids (colored peaks) with layers of density heatmaps representing different features, including class I (red) and II (blue) TEs, and meiotic ACRs (gray). The horizontal grey line represents the PCH. D) Normalized enrichment of ATAC-seq read coverage over repetitive elements of meiotic and somatic cells. E) Total coverage of ACR per region F) Total ACR coverage per genome feature. Upstream and downstream covers 1 kb from the transcription start and termination sites, respectively.

The enrichment of COs correlates with lower nucleosome occupancy and reduced DNA methylation ^31^. To investigate the chromatin state of TE elements with and without COs, we performed an ATAC-seq analysis of *S. lycopersicum* meiotic and somatic cells and found 52,802 and 25,101 accessible chromatin regions (ACRs), respectively. These ACRs have an average size of 733bp and represent accessible chromatin in the *S. lycopersicum* parent. We performed Pearson correlation analysis of the read distribution over the genome which showed high similarity between biological replicates (**Supplementary Figure 14**). Based on a permutation test, we found significant overlap between COs and meiotic ACRs (z-score = 87.2), confirming the reports that COs occur in regions accessible to recombination machinery. In **Figure 3e**, we can see that crossover regions have higher ACR coverage compared to random genomic regions. Upon comparing meiocyte ACRs with TEs, we found that TE superfamilies enriched with COs have an accessible chromatin segments, whereas retrotransposons like *Gypsy, Copia* and *L1* are not associated with accessible chromatin (**Figure 3d**). This is similar to the reports in *A. thaliana* of DNA transposons showing nucleosome depletion and higher SPO11-1-oligo levels ^31^. Moreover, retroelements like *Gypsy, Copia* and *L1* have very few SPO11-1-oligos, and high DNA methylation and nucleosome occupancy. This association of CO with specific class I and class II TEs is also observable in the landscape in **Figure 3c,** where the former are densely distributed in the PCH while the latter are predominantly found in DEU. Furthermore, the chromatin accessibility of TE superfamilies flips between the somatic and meiotic cells, hinting at a preference of keeping specific superfamilies inaccessible during meiosis (**Figure 3d**). The differential ACRs suggests that TE superfamilies may have different roles or activities between tissue types and in relation to recombination. Our results emphasize the major role of chromatin structure in the suppression or enrichment of COs in TEs and the need to particularly analyze meiocytes to account for tissue-specific ACRs.

Similar to the association of COs with proximal promoter regions ^23^, it was previously reported that ACRs are strongly associated with transcription start site (TSS) ^43^. To confirm it, we examined the average profile of ATAQ-seq signal in genes and their flanking regions, and found the highest coverage at TSS for both meiotic and somatic cells (**Supplementary Figure 15**). We also checked percentage of ACRs in genome features and discovered that the majority are located near or within genes (**Figure 3f**), similar to COs (**Figure 1a**). Normalized by the total genome coverage of the feature, the promoter regions and the UTRs (untranslated regions) have the highest ACR density.

Aside from the fact that many SVs are generated by *Gypsy* and *Copia* retrotransposons ^34^, 67% of coldspots we detected are covered by these TE elements for at least 50%. About 98.6% of the conserved coldspots are in PCH where retrotransposon presence is dense ^31^. Furthermore, the retrotransposon families that are linked with CO suppression, cover 450Mb (~52%) of the tomato genome, implying the wide span of suppression due to retrotransposons. This underscores the importance of transposable elements in shaping recombination patterns, both in hybrids and inbreeding materials and predominantly in regions with high retrotransposon density.

### Supergenes and breeding bottlenecks

COs tend to occur near genes but certain genomic elements prohibit the recombination between loci, causing co-segregation of these loci to the offspring. About 62% of genes are in CO coldspots of one or more of the hybrids and 484 of these coldspots contain at least 20 genes (**Figure 4a**). Gene complexes within coldspots are supergenes in a specific hybrid, but the same supergenes are not necessarily found in other hybrid tomato crosses. Although many supergenes are located in the conserved coldspots in PCH, other supergenes are located in the 81 coldspots in gene-dense DEU. In a gene ontology (GO) enrichment analysis of crossover and coldspot regions (**Figure 4b**), we found that 45 biological processes, 14 molecular functions, and 48 cellular components are significantly enriched (false discovery rate < 0.05) and that overrepresented GO terms in coldspots are associated with basal housekeeping functions (e.g like transcription coregulator activity, transporter complex, rRNA processing, metabolic processes). A more detailed list of enriched GO terms is reported in **Supplementary Figure 16**. Interestingly, we identified multiple metabolic processes enriched in the coldspots (**Supplementary Figure 16a**), which may reflect the evolutionary divergence between the tomato and wild parents. Many of the coldspot genes are directly related to modification in metabolism, which is considered a prominent manifestation of the domestication process ^44–47^. Supergenes, which are mostly generated by inversions or translocations, have actually been linked to metabolic pathways and alternative phenotypes in plants ^11,45^.

**Figure 4.**
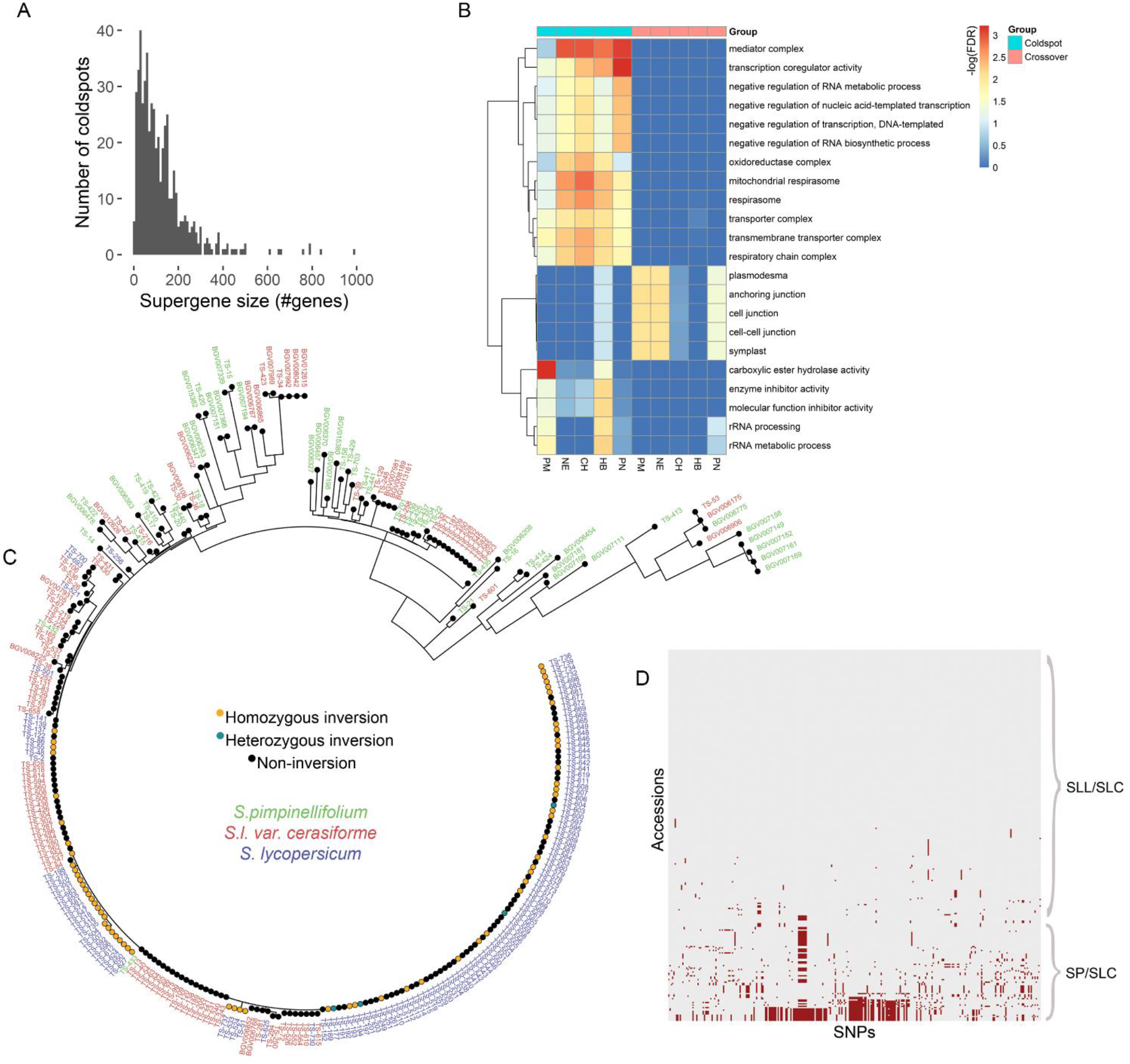
Supergenes and inversions. A) Sizes of supergene clusters in CO coldspots. B). Gene ontology (GO) terms enriched (at least 2x) in coldspot and crossover regions. C) Phylogenetic tree of genic SNPs in an inversion region of wild and domesticated tomato accessions. D) SNPs w.r.t *S. lycopersicum* within the inversion region.

To further investigate links between coldspots and phenotypes, we characterized the supergenes residing in the large coldspots in PN (**Figure 2e**). These coldspots contain 2,736 genes, 877 of which have been identified as domestication syndrome genes ^48^. The coldspots in the short arms of chromosome 6 and 7, coinciding with the inversion as previously reported by Szinay, et al. ^9^, contain 130 and 295 genes, respectively, and are associated with responses to oxidative stress (P = 1.27 x 10^-4^) and specific catabolic and metabolic processes (9.66 x 10^-8^) (**Supplementary Figure 16**). Similarly, the coldspots in both arms of chromosome 1 contain genes involved in metabolic processes of organic substances (P = 2.26 x 10^-5^) and transcription coregulator activity (P = 6.06 x 10^-5^). This result does not simply imply association between metabolic processes and coldspots, but it underlines the highly constrained metabolic profile in hybrids or in tomato introgressed with these coldspot regions.

Aside from the rewiring of the metabolome, we are interested in knowing if domestication is associated with linkage drag in regions containing resistance (*R*) genes. Upon inspecting the coldspots in PN, we found that they are enriched with *R* genes (Fisher’s exact test; P = 5.1 x 10^-4^) and at least 29 coldspots (23 clusters) contain *R*-gene hotspots (**Supplementary Figure 17**). The coldspot in chromosome 7 contains 295 genes, including *R* genes and *chitinase* genes. In this region, we found an enrichment of genes related to the *chitin catabolic process* (FDR = 1.49 x 10^-4^), *chitin binding* (FDR = 1.63 x 10^-4^) and *chitinase activity* (FDR = 2.18 x 10^-2^), which are involved in plant defense responses against pathogens ^49–51^. Our findings is consistent with the observation in Arabidopsis, in which CO coldspots with many SVs contain clusters of *R* genes ^52^. Some *R*-gene hotspots can become CO hotspots to overcome new pathogens through rapid diversification. In contrast, *R* genes conferring resistance to pathogens with low genetic plasticity are located in CO coldspot, possibly maintained by the structural heterozygosity ^53,54^. The association between resistance genes and some unfavorable alleles due to genetic linkage limits the introgression of resistance haplotypes into breeding lines. A specific case of linkage drag involving the resistance to *Fusarium* wilt race 3, reduced fruit size and increased sensitivity to bacterial spot, was broken by reducing the size of the introgression ^55^. However, this shrinking of introgressed region is feasible only because it is not induced by an inversion or other CO-suppressing type of SV, unlike the *Ty-1* and *Ty-2* introgression which both are located within inversions ^20^. Aside from *R* genes, the coldspot in chromosome 7 also contain the *SUN* locus, which is linked with variable fruit shape in the wild and cultivated tomato ^56,57^. The remaining coldspots in PN contain 17 genes with putative roles in fruit shape determination, further substantiating the association between coldspots and domestication syndrome traits ^58^.

To break linkage drag in an SV region with no recombination, alternative crosses that do not result to coldspot is needed. For the coldspot in chromosome 7 of PN, the other wild relatives can serve as alternative parent as they all exhibit recombination in this region. Another example of region with no recombination is a 294-kb euchromatic inversion in the *fasciated* (fas) locus with breakpoints in the first intron of a YABBY transcription factor gene (*SlYABBY2b*) and 1 kb upstream of the *SlCV3* start codon ^59–61^. This inversion contains 41 genes, including 4 disease resistance genes, and knocks down YABBY, conferring a large fruit phenotype to domesticated tomato. Given the resequencing data for populations of wild and domesticated tomato, it might be possible to find “bridge accessions” or accessions without the allele causing heterozygosity. We therefore screened 56 accessions of wild (*S. pimpinellifolium;* SP), 109 early-domesticated (*S. lycopersicum* var. *cerasiforme*; SLC) and 127 vintage tomato (*S. lycopersicum* var. *lycopersicum*; SLL) that were genotyped for the inversion, including the SNPs within the inversion. All SP and 96% (109) SLC accessions have non-inversion genotypes while half (64) of the SLL group have at least one inversion allele (**Figure 4c**), which may suggest that the inversion could have occurred during tomato domestication. This is consistent with the drastic reduction of nucleotide diversity in this region when comparing SLC/SLL with SP population ^48^. Upon inventorying the inversion and non-inversion accessions, we compared the SNP profile within the inversion region and found distinct haplotypes of SP compared to the SLL accessions (**Figure 4d**). We subsequently identified at least 12 SLL accessions without the inversion and with larger fruit weight phenotype compared to SP ^48^. These candidate bridge accessions may be crossed with SP accessions to overcome CO suppression in the inversion region while maintaining genetic background that confers large fruit other than the *fas* inversion. Aside from this inversion, there are at least 236 additional non-overlapping SV sites in the population of SP and SLC/SLL that may be analyzed to predict recombination barriers, especially lineage-specific rearrangements. Most importantly, this SV profile may be used to select for bridge accessions to introgress genetic diversity into genetically eroded domains of crop tomato.

## Discussion

COs are mostly distributed in the gene-rich DEU regions of each chromosome, consistent with previous reports ^22,23^. The recombination landscape in one of the hybrid accurately matched a genetic linkage map, underscoring the importance of our method which provide high resolution CO with less cost and labor. Despite the similar overall CO landscape between hybrids, we discovered fine-scale differences in CO patterns and regions without recombination. CO coldspots have limited ability to reshuffle alleles between tomato and wild species, which hamper introgressive hybridization breeding and reduce efficiency of backcrossing. Although the majority of the coldspots are conserved between all hybrids, some coldspots are unique to a cross, which may serve as putative targets to break linkage drag or to study the underlying fitness advantage that necessitates the suppression of recombination. This is so far the most comprehensive profile of recombination in tomato hybrids and possibly even in plant species.

Across all hybrids, we found conspicuous absence of crossover events in SV regions, particularly in lineage-specific rearrangements. The varying patterns of recombination between the hybrids is associated with the rearrangements between the wild parental genomes, implying that SV profiles in F1 progeny may help distinguish regions that may or may not allow crossovers. Knowing these regions enables breeders to fine-tune introgression plan by inspecting recombination patterns in the loci of interest prior to the elaborate hybridization and screening processes. Although multiple studies have already reported the negative association between SV and COs, it is still not clear how SVs inhibit recombination. Rowan, et al. ^6^ proposed several possible explanations for the observed suppression of COs in heterozygous SVs, such as absence of a repair template, tendency to produce non-viable gametes, DNA methylation in the SV region, and blocking physical interaction in variant regions preventing proper synapsis. Furthermore, it has been reported that DSBs in inversion regions are preferentially resolved as noncrossover gene conversions and not as COs ^6,62,64^. Although we have already found a association between CO and SV patterns, further studies must be conducted to improve the detection of SVs, specifically of the insertion and translocation type, for better recognition of the underlying causes of suppression in each coldspot.

Structurally heterozygous regions in the genome, that cause lack of recombinant haplotypes, have been linked to adaptive phenotypes and plant domestication and speciation ^11,12,15,38^. An inversion, capturing two or more alleles adapted to an environment, prevents recombination and confers a selective advantage that subsequently promotes its spread in the population ^64^. Based on visual examination of the synaptonemal complexes, it was suggested that SVs form interspecific reproductive barriers in the tomato clade ^21^. We confirmed it by our results on the absence of COs in SV regions of multiple interspecific crosses. Some of these recombination coldspots contain supergenes which may confer alternative or differentiated phenotype between the parental genomes ^11^. This information can help identify unfavorable gene complexes prior to the hybridization, exposing possible undesired consequences of introgression. In this study, we showed that some CO coldspots in interspecific hybrids overlap *R* gene hotspots, which not only accumulated nucleotide variations during the evolution of wild tomato relatives but underwent copy expansion and contraction, conferring varying resistance to pathogens ^65^. But the CO suppression prevents traditional introgression methods from selecting favorable alleles and gene copies in these *R* gene hotspots ^66^, curbing the efforts to develop disease-resistant tomato. Furthermore, some coldspots contain genes associated with metabolic processes and fruit traits, implying linkage between genes that may relate with the considerable change in chemical composition of tomato fruit due to fruit mass-targeted selection during domestication ^47^. These coldspots can serve as targets for metabolite engineering in *de novo* domestication of wild tomato relatives. The enrichment of genes in CO coldspots linked with resistance and metabolomes is partly brought about by plant evolutionary events involving SVs ^38,45^. Further examination of recombination coldspots can help breeders to understand the genetic or epigenetic cause of CO suppression and determine divergent phenotypes resulting from the evolution of locally adapted alleles and from domestication.

Aside from the association between SVs and CO coldspots, we also found specific superfamilies of TEs exhibiting strong association with crossovers and accessible chromatin regions. By checking the ACRs in meiocytes, we determined that the varying association between superfamilies may be influenced by their chromatin configuration, keeping elements like *Gypsy* and *Copia* inaccessible during meiosis which consequently prohibits COs. We also discovered differential chromatin accessibility of TE elements in somatic and meiotic cells, necessitating further studies to explain whether this relate with different functions or the regulation to limit proliferation of specific TEs during meiosis ^67^. Although we found an association between TEs and COs, it is not clear whether TEs directly shape the recombination landscape, or that recombination and TE insertions simply collocate in ACRs and genic regions because of TE insertion bias ^31,40^. In tomato, *Stowaway* elements preferentially inserts within or near genes while *Gypsy* elements inserts in pericentromeric regions ^68,69^, agreeing with the correlation of COs and TEs. On the other hand, the consistent chromatin state per TE superfamilies may indicate that, depending on the type, new TE insertions can either suppress or promote recombination ^31,40^. For example, the expansion of pericentromeric regions in *A. alpina* due to retrotransposon insertions resulted to more regions with suppressed recombination^70^. It would also be interesting to further examine how the activity of TEs, such as during stress exposure, can influence the recombination landscape ^71,72^. With the findings we have, the value of both SV and TE profiles in the parental genomes for CO hotspots and coldspots prediction becomes more apparent.

Previous studies have actually tried to increase CO frequencies but failed to do it homogeneously along the genome ^73–75^, missing regions CO coldspots. Recently, it was demonstrated that recombination can be restored by inverting an inversion using genome editing (Schmidt *et al*., 2020). However, current regulations may restrict how solution to break linkage drag based on genome editing may serve direct breeding applications. Alternatively, we demonstrated that we can find bridge accessions that can solve the lack of recombination in regions with SVs while maintaining the desired genetic background. However, it is dependent on whether those accessions exist in nature or not and on the comprehensiveness of the resequencing data. Recent work by Alonge, et al. ^34^ involves the profiling of SVs in 100 subset accessions that represent the diversity of over 800 tomato accessions, providing more data that can be used in finding compatible genomes. If we cannot find a bridge accession and the SV of interest is heterozygous in one of the parents, it is possible to screen for a homozygous genotype from an offspring population. Nevertheless, we emphasize the importance and advantage of doing compatibility or linkage drag checks, in a cost-effective way, as part of the breeding scheme. Future work can focus on profiling CO-associated features in resequencing data of tomato and wild relative populations and on predicting CO coldspots between a pair of accessions without developing a mapping population.

## Methods

### Sequencing of pollen gametes

We produced F1 plants from crosses between *S. lycopersicum* cv. Heinz1706 and the following wild relatives: *S. pimpinellifolium* (CGN14498)*, S. neorickii* (LA0735), *S. chmielewskii* (LA2663)*, S.habrochaites* (LYC4), and *S. pennelli* (LA0716). The wild species served as the male parents. Mature pollen were collected from each hybrid and processed to isolate the high molecular weight DNA using the protocol in Fuentes, et al. ^23^. 10X Genomics libraries were constructed according to the Chromium™ Genome v2 Protocol (CG00043) and then sequenced on an Illumina HiSeq 2500. Aside from the sequencing pool of pollen from these hybrids, we used the same protocols to sequence the inbreds of the parental tomato and the wild species.

### Crossover detection

For detecting segregating markers in the hybrids, linked reads from the inbred wild parents were aligned against the *S. lycopersicum* cv. Heinz reference genome SL4.0; ^76^ using *Longranger* ^77^ and were subsequently processed using GATK HaplotypeCaller ^78^ with the recommended hard filtering to screen single nucleotide polymorphisms (SNPs). Heterozygous SNPs and other SNPs located in homopolymeric regions and regions prone to false positives due to inaccurate assembly or copy number variations, resulting in highly heterozygous alignments, were filtered out. Thereafter, for each hybrid, the linked reads from pollen gametes were aligned against SL4.0 using *Longranger* and were phased using the segregating markers as described in Fuentes, et al. ^23^. For each putative recombinant molecule, we applied filters on the resolution, spanning distance, block size, and the number of supporting reads, wild cards and markers per phased block. In the updated version of our pipeline ^23^, filtering putative recombinant molecules with significant overlap with repeats and transposable elements was deprecated to enable analysis of correlation between COs and superfamilies of TEs. The number of overlapping crossover events between hybrids and their significance were determined using *bedtools* ^79^. To compare the landscape, Pearson’s correlation matrix was computed on the CO count in 500-kb windows with 50-kb step size.

### Detection of coldspots

We counted the number of COs per hybrid in 10kb sliding windows and merged those windows with at least one CO and within 1kb distance of each other. The resulting set of genomic intervals are considered CO regions. Regions without COs spanning at least 1Mb are considered coldspots. To cluster coldspots from all hybrids, we first grouped those with at least 1 bp overlap. For each group, we built a graph with coldspots as nodes, connected by edges if they have a least 50% reciprocal overlap. Each graph was split into connected components (C) and then based on the genomic position, we computed the distance (*p*_k_) between the leftmost and rightmost coldspot in each component. If *p*_k_ is at least 1.5 times the size of the smallest coldspots in C_k_, the component was further regrouped by hierarchical clustering using a distance matrix d(i,j) = (*f*-2*length(i∩j))/(*f*-length(i∩j)), where i and j is the pair of coldspots in a component and *f* is the sum of their lengths. Hierarchical clustering by complete linkage was used; the resulting dendrogram was cut at the height of 0.3. The resulting groups were used to define shared coldspots, which occur in at least two hybrids, and unique coldspots *i.e*. those coldspots that occur only in one hybrid. We also identified conserved coldspots or regions without CO in all five hybrids.

### Detection and validation of SVs

Linked reads from inbreds of all the parental species were aligned to the reference genome and analyzed to detect SVs using *Longranger*. With the presence of heterozygous SVs in the parental genomes, it is possible that only the reference allele may have been inherited by the F1 plants. To determine for each hybrid whether the F1 plants inherited an SV allele causing heterozygosity between the homologous chromosomes during meiosis, we profiled SVs in the F1 pollen linked-reads. The pool of pollen included both recombinant and non-recombinant regions and represented alleles from both parental genomes of each F1 plant. Thereafter, SVs were reported if present in both the inbred and the corresponding pollen data, referring to them as *parental SVs*. To further remove problematic regions, SVs between the Heinz reference and the Heinz inbred, which we refer here as *self SV*, were detected. Lastly, we reported parental SVs, of the deletion (DEL) and inversion (INV) type, that do not overlap self SV. For SV validation, we compared the SL4.0 assembly against the existing assembly of *S. pennellii* ^80^ using Syri ^81^ and manually inspected randomly selected sets of DELs and INVs using Gepard ^82^.

### Enrichment analysis

To determine the enrichment of COs in specific TE superfamilies, we generated 10,000 permutations of the CO data per hybrid using bedtools and computed the number of overlaps with transposable elements. We then compared the observed and the expected overlap with TE of these CO regions. For detecting overrepresented motifs, we retrieved the genomic sequences spanning CO regions with a resolution above 0.002, including the 3-kb flanking regions, and analyzed these with the MEME suite ^83^ using default parameters. Furthermore, we generated a list of genes present in the CO and coldspot regions and subsequently ran Panther ^84^ to identify enriched GO Terms. We also computed the number of resistance genes ^53^ and historical recombination hotspots ^15^ in the CO coldspots.

### Genotyping of population data

Using *bwa mem* ^85^, we aligned a set of resequencing data for 357 accessions compiled in Fuentes, et al. ^15^ against the SL4.0 reference genome. SNPs were detected using GATK HaplotypeCaller and were further filtered using GATK joint-genotyping and hard filtering. We then selected biallelic SNPs with a minimum allele frequency of 0.05 and less than 10% missing data using bcftools ^86^ and imputed missing calls using Beagle v 5.1 ^87^. To detect SVs, we ran Delly ^88^ for each accession and then we genotype SV sites across all accessions. We selected an inversion event and filtered accessions with missing calls, retaining 292 accessions. With SNPs in this inversion region, we generated and visualized the neighbor-joining tree using Mega7 ^89^ and Figtree (http://tree.bio.ed.ac.uk/software/figtree/), respectively.

### ACR Detection

Tomato plants were grown and cultivated in a greenhouse with a photoperiod of 16 hours light and 8 hours dark, and a minimum temperature of 16°C. Only healthy four- to seven-week-old plants were used in all experiments. The youngest leaves (the most apical) were used to isolate somatic nuclei. Meiocytes were isolated from young flower buds containing anthers that were less than 2 mm in size. Microscopic analysis revealed that at this stage in anther development nearly all meiocytes are in prophase I.

For nuclei isolation, approximately 0.4 g of young tomato leaves, or anthers from 20 prophase I flower buds were collected and immediately chopped in 2mL pre-chilled lysis buffer (15mM Tris-HCl pH7.5, 20mM NaCl, 80mM KCl, 0.5mM spermine, 5mM 2-mercaptoethanol, 0.2% Triton X-100) until a homogenous suspension was obtained. The suspensions were filtered twice through Miracloth and subsequently loaded gently on the surface of 2mL dense sucrose buffer (20mM Tris-HCl pH 8.0, 2mM MgCl2, 2mM EDTA, 25mM 2-Mercaptoethanol, 1.7M sucrose, 0.2% Triton X-100) in a 15mL Falcon tube. The nuclei were centrifuged at 2200g at 4°C for 20 minutes and the pellets were resuspended in 500μL pre-chilled lysis buffer.

Nuclei were kept on ice during the entire sorting procedure. Nuclei were first stained with 4,6-Diamidino-2-phenylindole (DAPI) and examined for integrity and purity using a Zeiss Axioskop2 microscope. Once the integrity and purity of nuclei was confirmed, nuclei were sorted in a BD FACS Aria III sorter. A total of 50,000 nuclei were sorted based on their size, shape and the intensity of the DAPI signal, which indicates the ploidy levels of the nuclei. 2n nuclei were sorted from young leaf samples, while 4n nuclei, corresponding to meiocytes, were sorted from anther samples. After sorting, nuclei were once more checked for integrity and purity under a microscope. Nuclei were transferred from sorting tubes to LoBind Eppendorf tubes and centrifuged at 1000g at 4°C for 10 min and then washed with Tris-Mg Buffer (10mM Tris-HCl pH 8.0, 5mM MgCl2).

Tn5 integration was performed as previously published ^90^ on purified nuclei using the Nextera Illumina kit (Illumina, FC 121 1031) at 37 °C for 30 min. After tagmentation (insertion of the sequencing adapter into accessible chromatin), the tagged DNA was purified with a Qiagen MinElute PCR purification kit. To generate an ATAC-seq library for sequencing, tagged fragments were amplified by two successive rounds of PCR. In the first round of PCR, the fragments were amplified by only 3 PCR cycles using the NEBNext High-Fidelity 2xPCR Master Mix and the Custom Nextera PCR Primer 1 and barcoded sets of Primer 2. Subsequently, 2.5 μL of the PCR amplified DNA was subjected to quantitative PCR to estimate the relative amount of successfully tagged DNA fragments and to determine the optimal number of amplification cycles for the second round of PCR. The latter was estimated by plotting fluorescence values against the number of cycles. The number of cycles required for the second PCR amplification equals the number of cycles that results in 25% of the maximum fluorescent intensity ^91^. ATAC-seq libraries generated were purified using AMPure XP beads (Beckman Coulter) and quantified using Qubit DNA high sensitivity assay in combination with Tapestation D1000 prior to sequencing.

Sequencing was carried out using an Illumina NextSeq 500. A snakemake analysis workflow (https://github.com/KoesGroup/Snakemake_ATAC_seq) was used for the analysis of the ATAC-seq dataset with the default parameters of the configuration files. Briefly, paired-end sequencing reads were trimmed to remove the Illumina adapter sequences using Trimmomatic 0.38 ^92^. Only reads with a quality score (Phred) above 30 were kept and mapped to the SL4.0 version of the tomato genome, tomato chloroplast genome and tomato mitochondrial genome using Bowtie2 ^93^. Only reads mapping to a unique position in the tomato genome were used for further analysis. Reads mapping to the tomato genome were then shifted to correspond to the real Tn5 binding location using the Deeptools alignmentSieve with the parameter “ –ATACshift”. ATAC peaks were called using the MACS2 algorithm ^94,95^.

Reads mapping uniquely to the transposable element annotation were counted using bedtools. Read counts were normalized by the total number of reads in the library and then grouped by the transposable element classes. Heatmaps and clustering was performed using the pheatmap package 1.0.8 (CRAN).

## Acknowledgements

This project was supported by the MEICOM Marie Sklodowska-Curie Innovative Training Network (ITN), H2020-MSCA-ITN-2017 Horizon 2020 Grant agreement number 765212 and by the Netherlands Top Consortium for Knowledge and Innovation (TKI project LWV19283). We also thank ENZA Zaden Research & Development B.V., Bejo Zaden B.V., Hortigenetics Research (S.E. Asia) Limited, Syngenta Seeds B.V., and KWS Saat SE & Co KGaA for support. The parental seed material was donated by the Centre for Genetic Resource, The Netherlands.

## Author Contributions

R.F., S.P., D.R. and A.D. designed and initiated the study. T.H., W.D., H.B., and E.L. generated the 10x Genomics sequencing data. R.N. performed the SNP calling. R.F. performed all analyses on crossovers. J.C., P.F. and M.S. generated and analyzed the ACRs. S.P., D.R., A.D.,S.T. and H.J. contributed to interpreting the data. R.F. wrote the paper with additional input from all others authors.

## Supplementary Figures

**Supplementary Figure 1.**
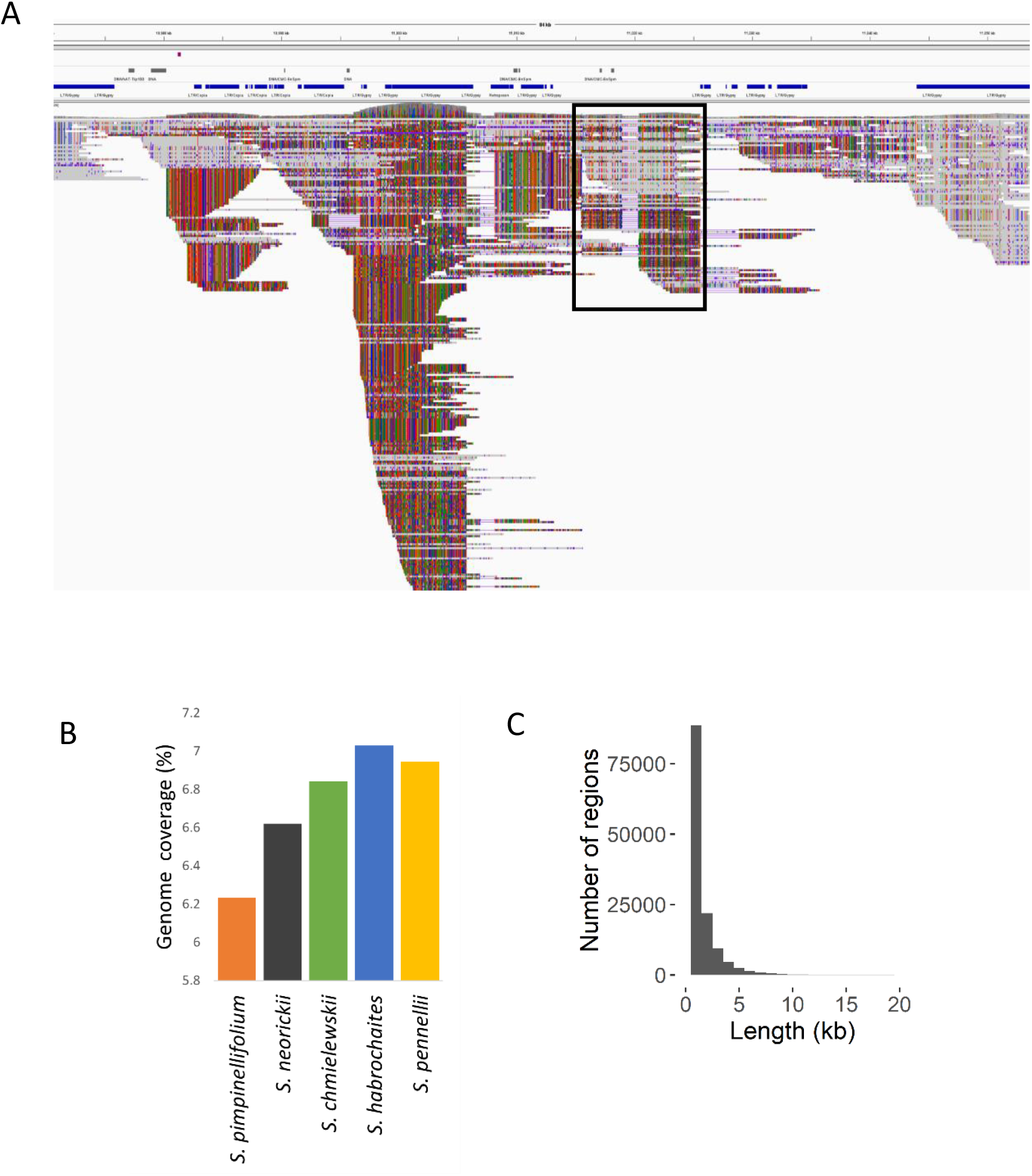
False positive hotspots in pericentric heterochromatin. A) Regions with excessive levels of heterozygosity and read coverage causing false positive crossovers (black box). Possibly, these regions are collapsed genomic segments in the reference genome or part of a copy number variation. B) The coverage and C) length distribution of these regions.

**Supplementary Figure 2.**
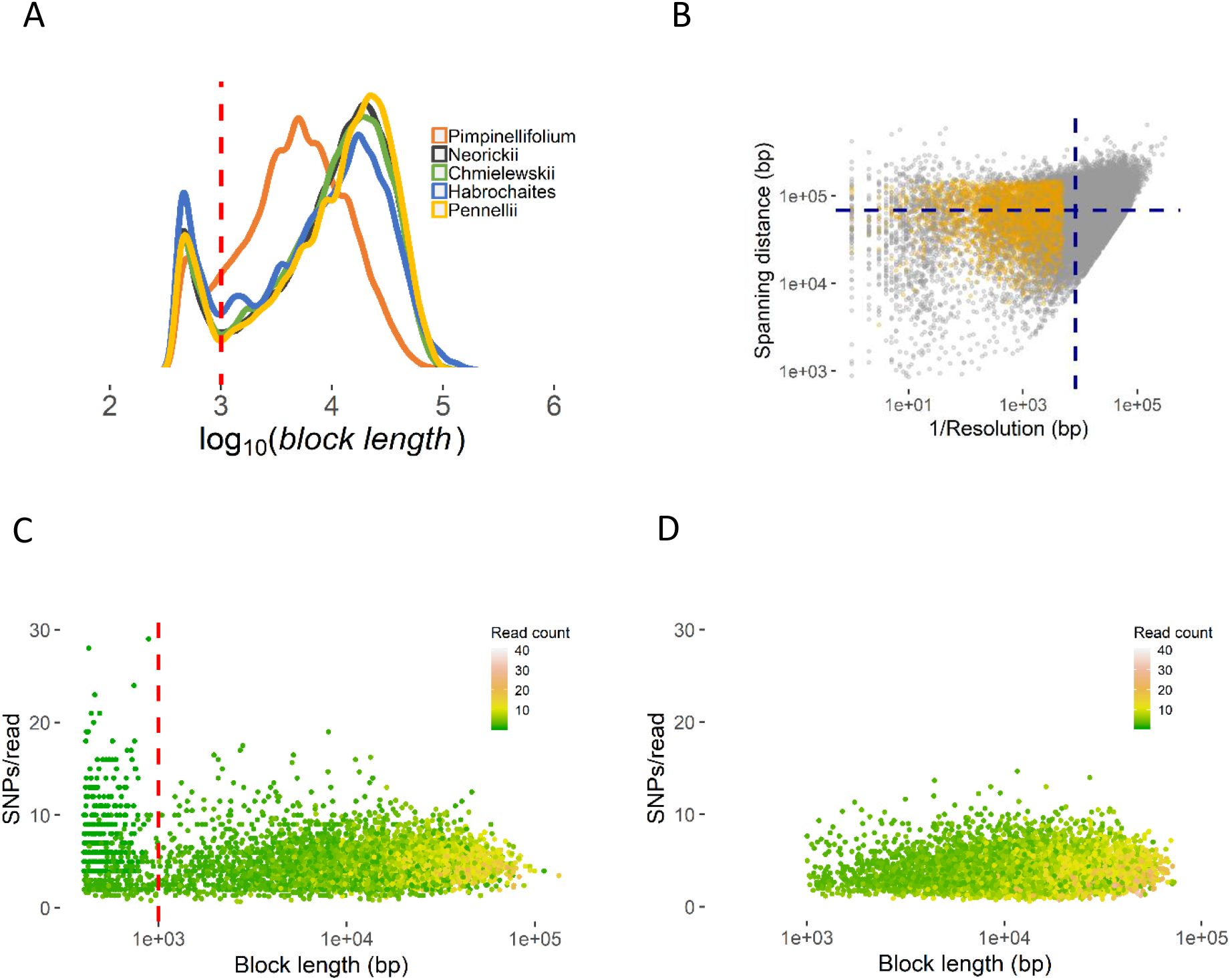
Filtering criteria on recombination molecules. A) A lower limit on the size of a haplotype block (marked by a red vertical line) is set based on the distribution of all block sizes across all crosses. B) Distribution of resolution and spanning distance of each recombinant molecule. The yellow dots represent COs that passed the filtering. The blue lines mark the average spanning distance and resolution. C) Ratio of SNP and read count per haplotype block as a function of block length. Haplotype blocks with sizes below 1kb are supported by few reads with high SNP density which may result from mismapped reads, further supporting the cut-off for haplotype block size. D) Final set of COs that passed all filtering constraints.

**Supplementary Figure 3.**
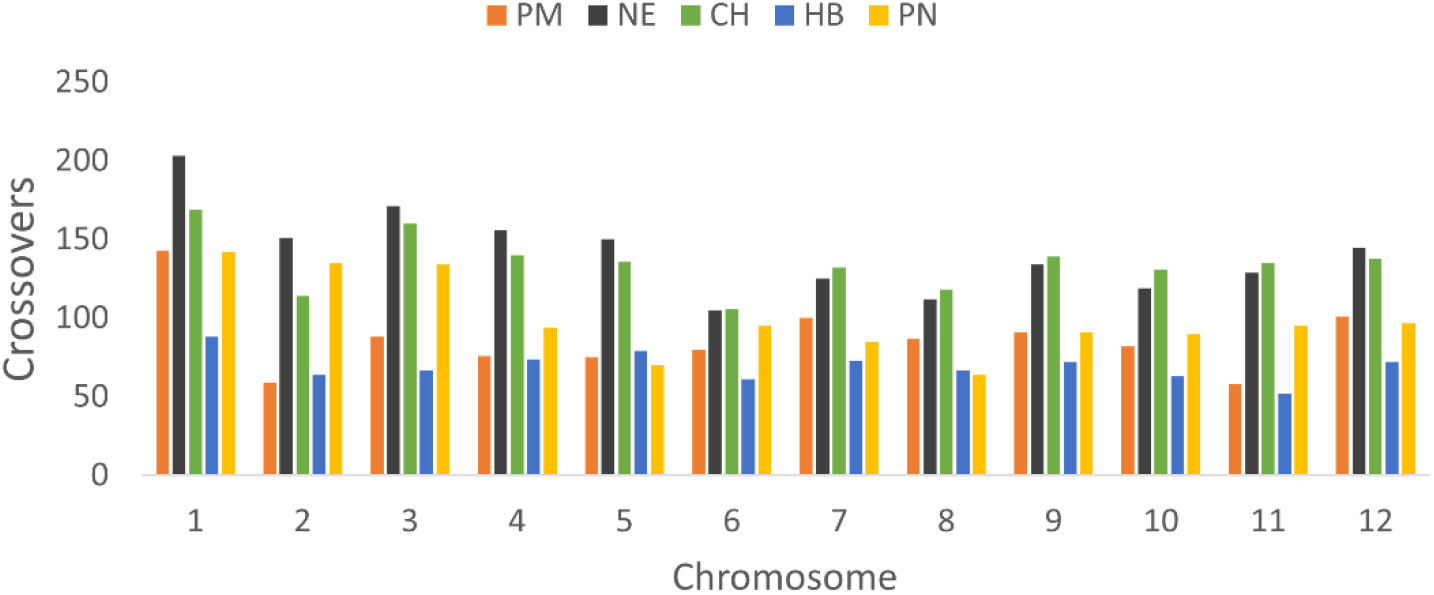
Frequency of crossovers per chromosome.

**Supplementary Figure 4.**
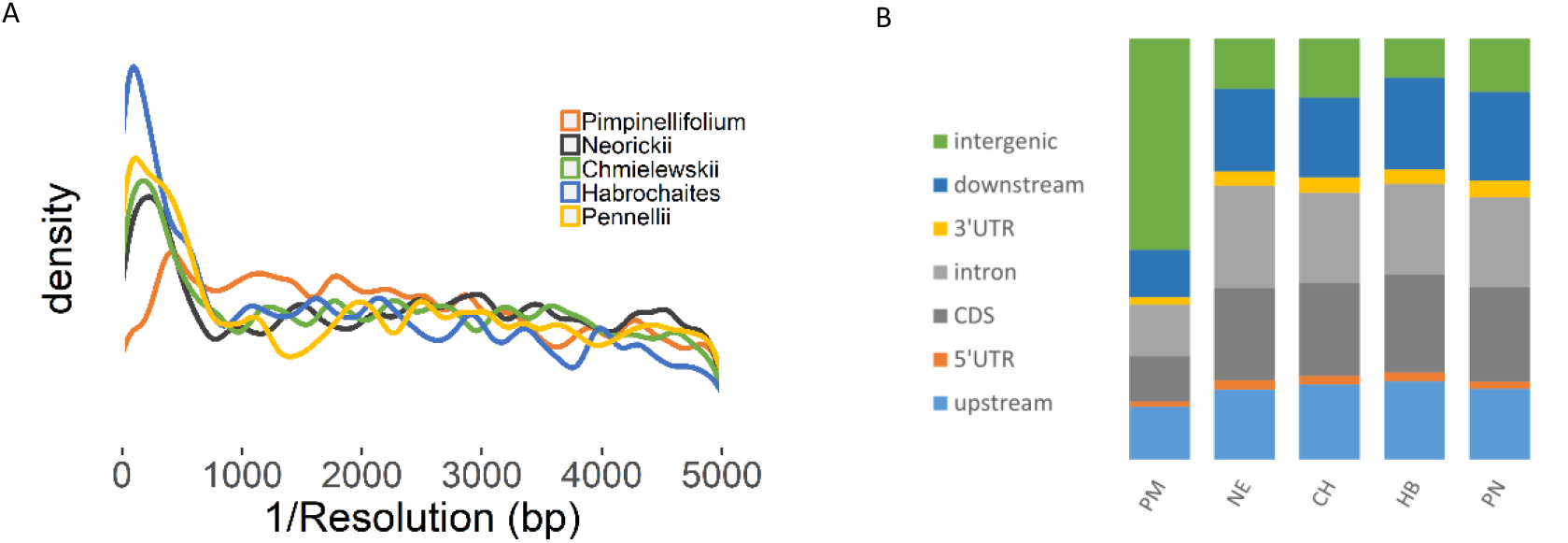
Crossover resolution and gene overlap. A) *S. pimpinellifolium* crossovers have lower resolution compared to the other groups, but between the resolution of 0.0002 to 0.001 (1kb to 5kb), the distributions are similar across the different crosses. B) Overlap of crossovers (resolution between 0.0002 to 0.001) with gene and intergenic regions.

**Supplementary Figure 5.**
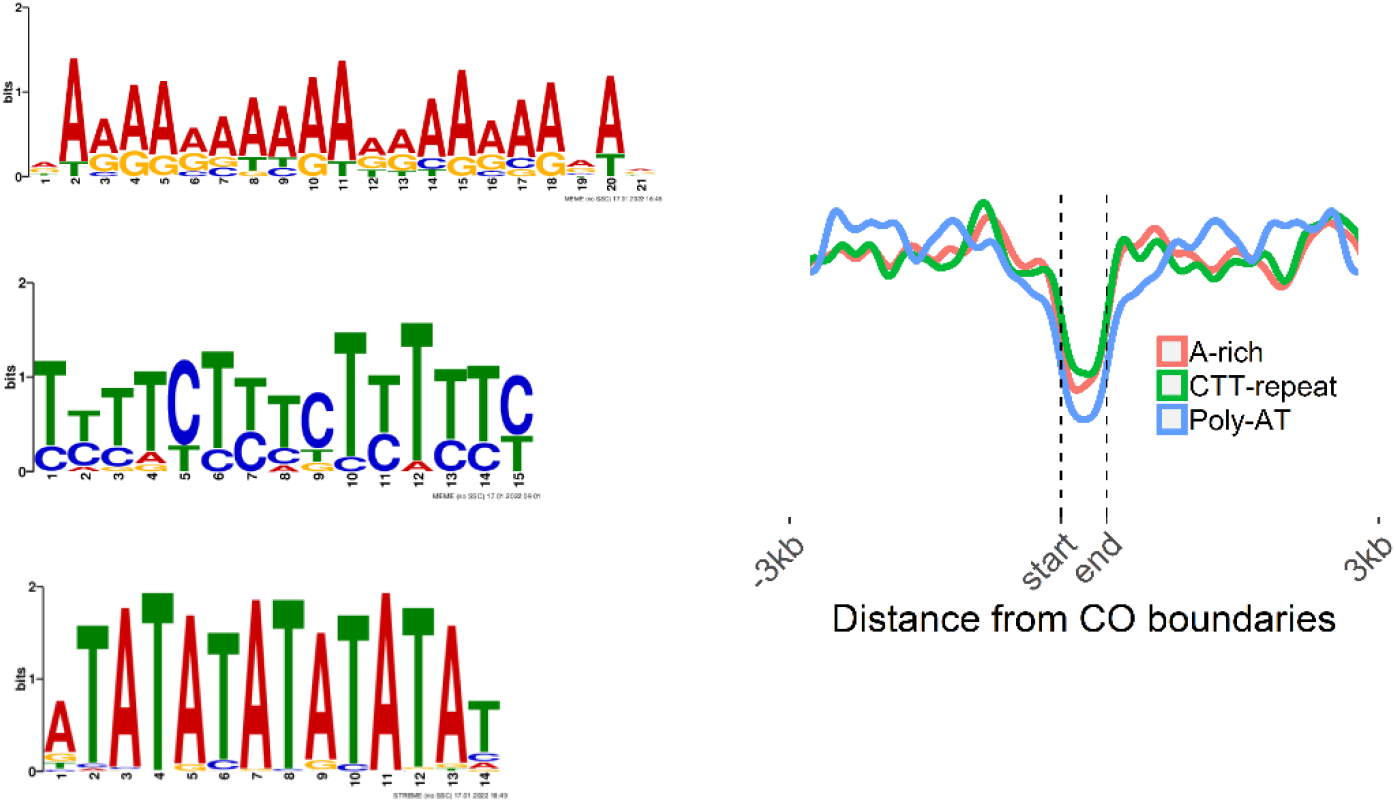
Overrepresented motifs. A) Motifs found within and flanking CO regions. Of 1,267 COs with resolution of at least 0.002, only 8-28% have the motif within the CO regions while the rest contain multiple copies of the motifs in the flanking regions. B) Distribution of the motifs within and around high-resolution COs.

**Supplementary Figure 6.**
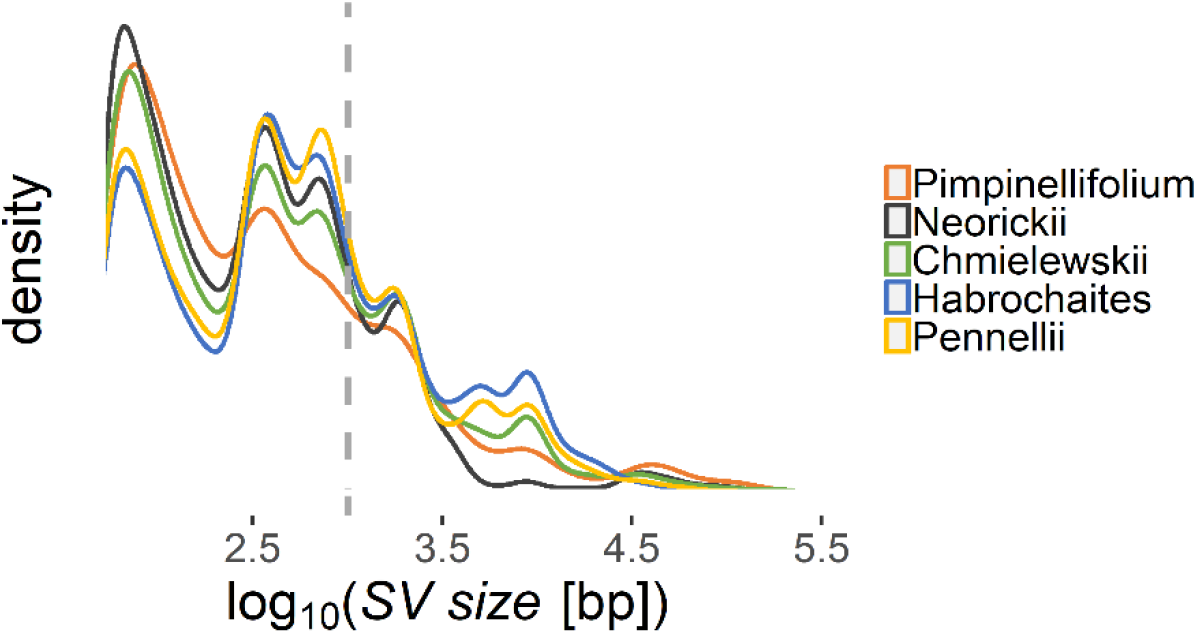
Longer structural variants for more distant wild genomes. Distribution of SV sizes per population showing higher frequency of longer SVs for *S. habrochaites* and *S. pennellii*.

**Supplementary Figure 7.**
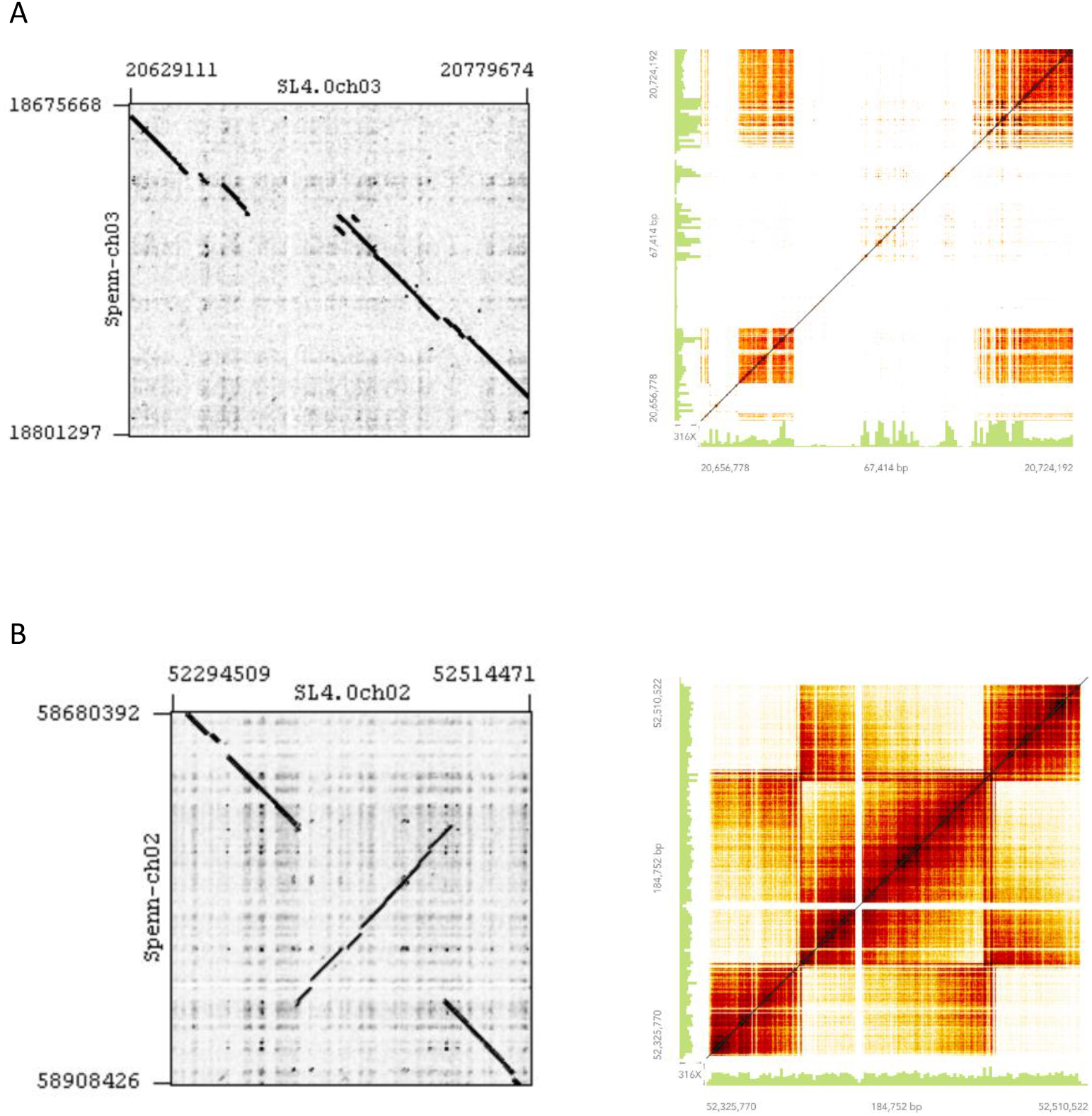
Validation of structural variants. Examples of a (A) deletion and an (B) inversion that are validated by manual inspection. First, through dot plots between the assemblies of *S. lycopersicum c.v*. Heinz 1706 and *S. pennellii* genomes generated using *Gepard* (left). Second, through heatmap of overlapping barcodes between linked reads (10X Genomics) in the *S. pennellii* parental genome generated using *Loupe Browser*. The patterns in the top right and bottom right figures characterize a deletion and an inversion, respectively.

**Supplementary Figure 8.**
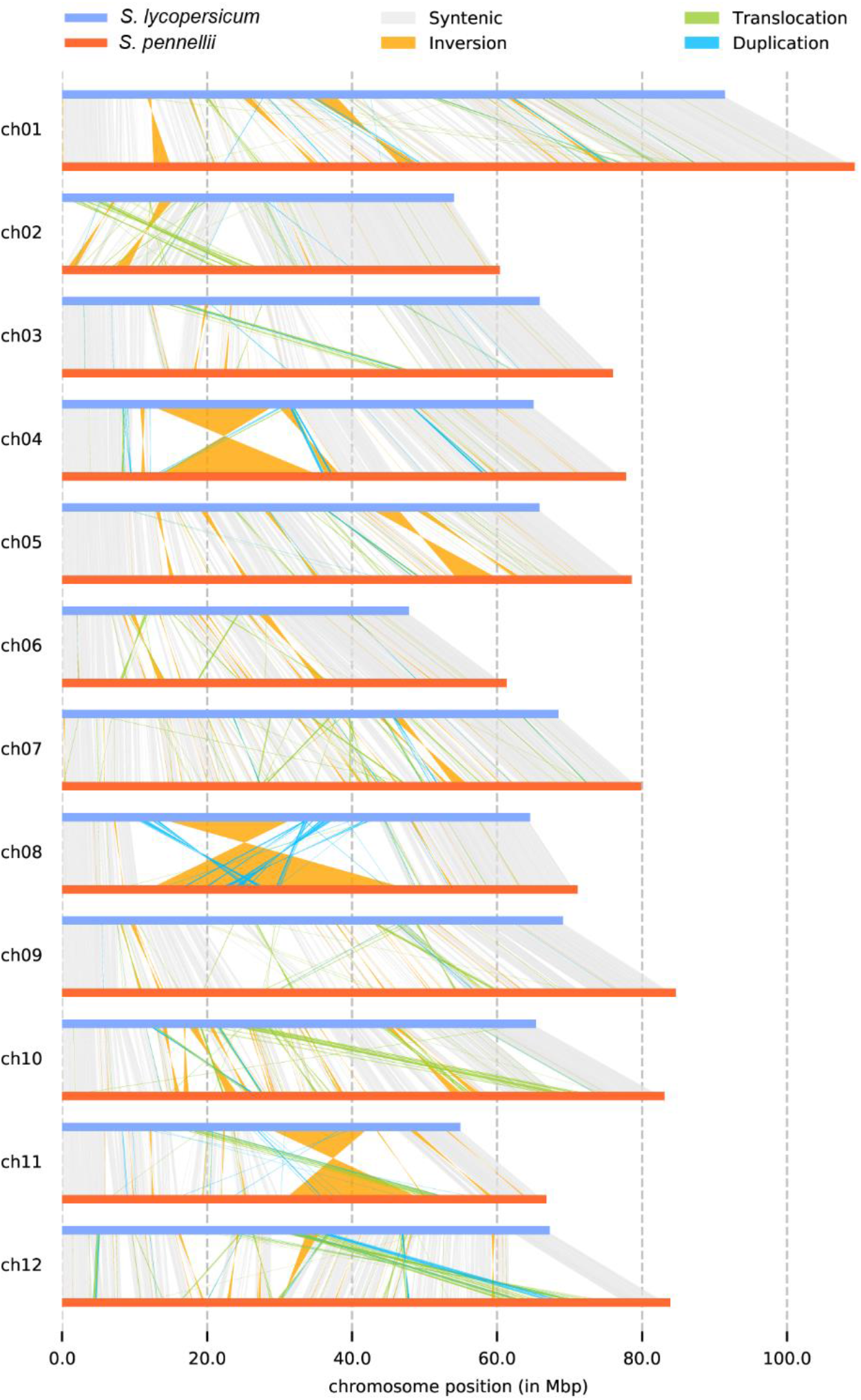
Parental genome alignment. Alignment between the assemblies of *S. lycopersicum* and *S. pennellii* showing syntenic regions and rearrangements.

**Supplementary Figure 9.**
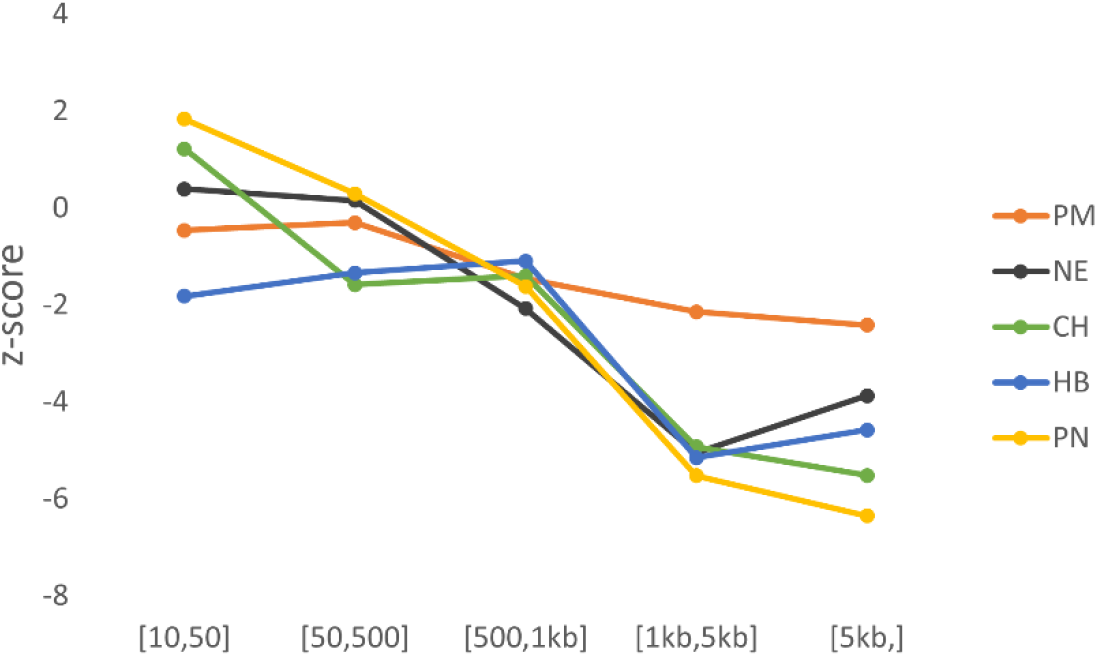
Suppression of COs relative to the size of SVs. To determine the relation between SV size and CO suppression, SVs are first binned according to size. Afterwards, per bin, the overlap of SVs and COs in the observed data was compared against the overlap in 10,000 permutation sets. Only bin [1kb,5kb] and [5kb,] have significantly fewer COs in SVs than expected by chance for all populations.

**Supplementary Figure 10.**
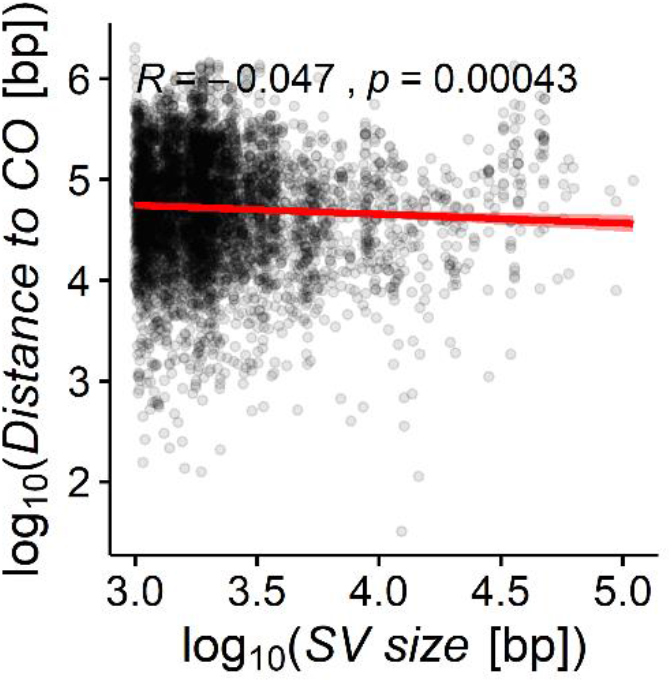
Distance of SVs to COs by size. There is no association between SV size and the distance of the nearest CO.

**Supplementary Figure 11.**
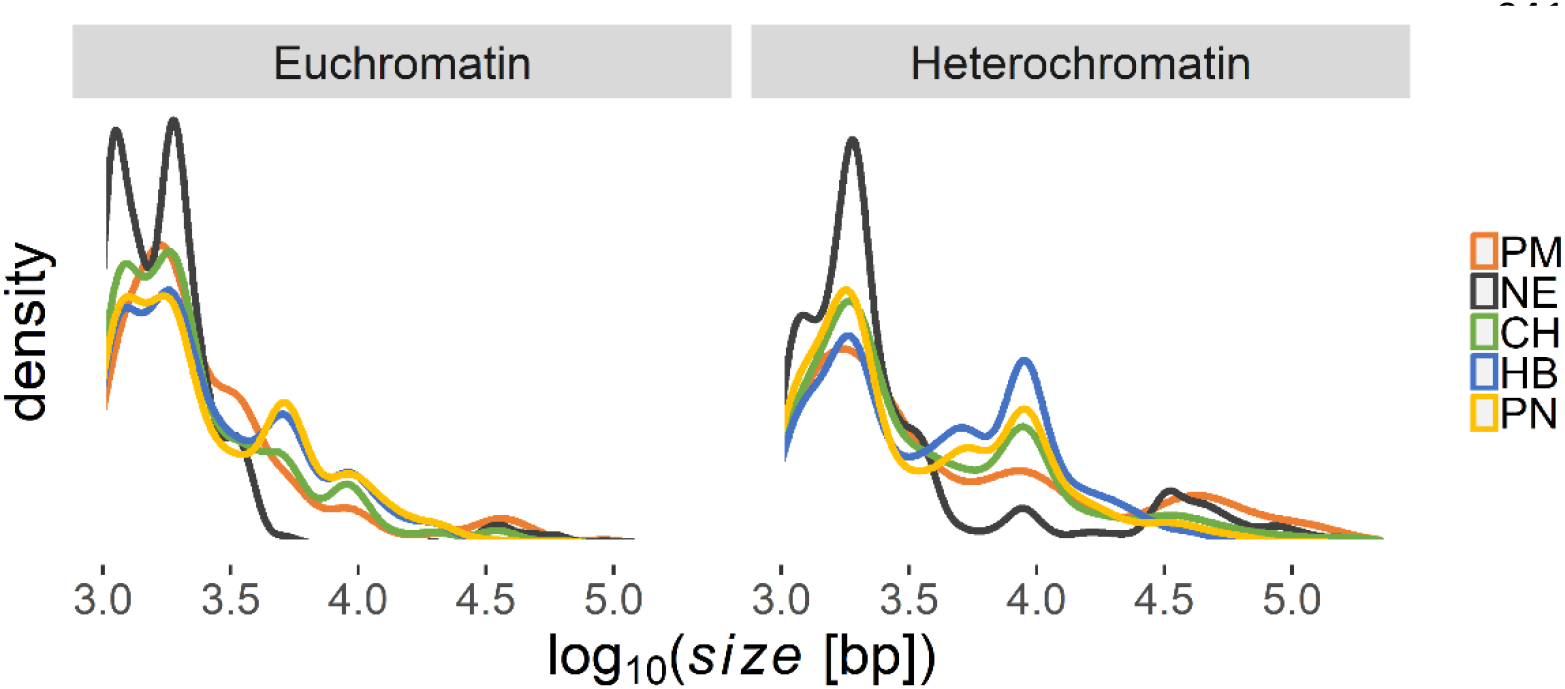
Sizes of SVs. Pericentric heterochromatin contains longer SVs in most wild genomes.

**Supplementary Figure 12.**
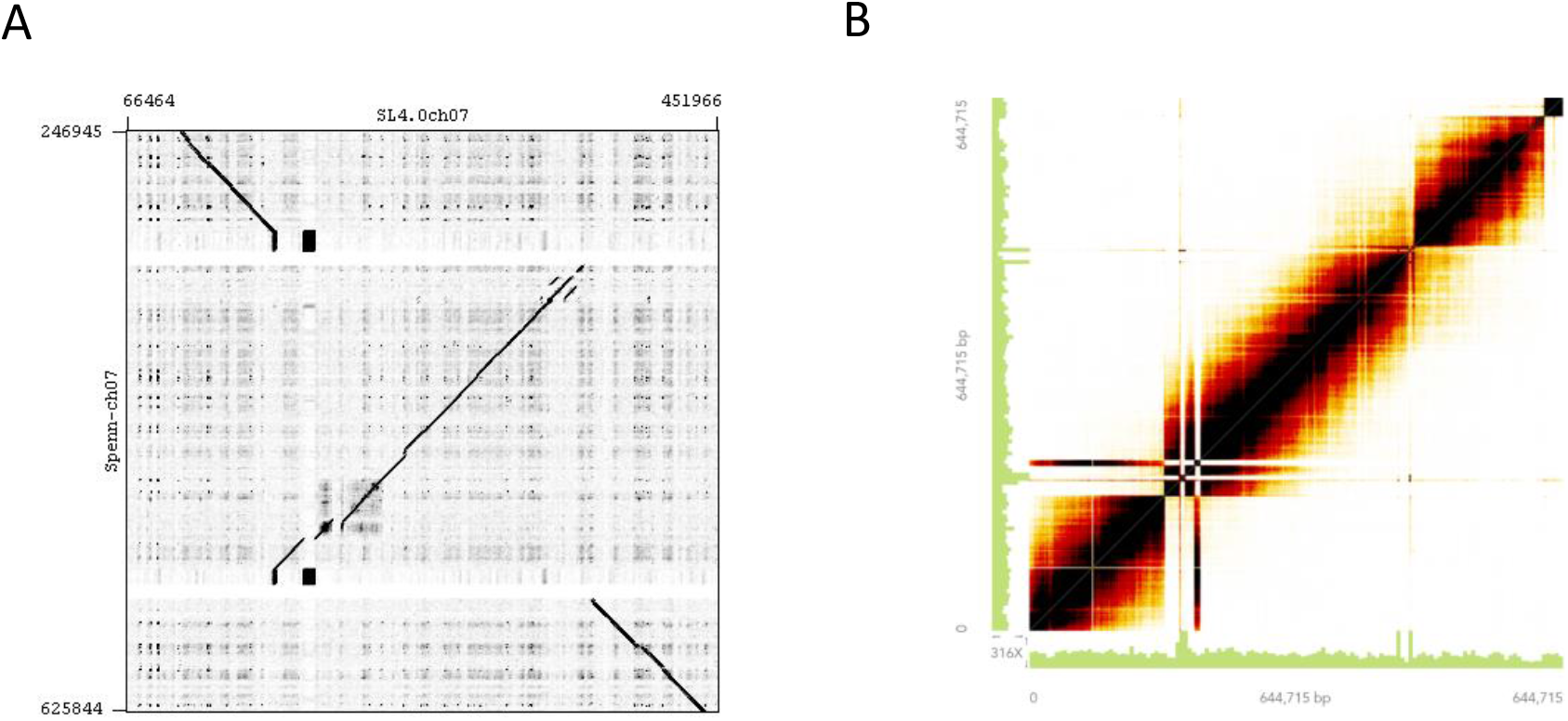
Inversion in chromosome 7 short arm. A) A distal inversion between the short arm of *S. lycopersicum c.v*. Heinz 1706 and the *S.pennellii* assembly was visualized using a dot plot. B) Heatmap of overlapping barcodes between linked reads (10X Genomics) in the inversion region of *S. pennellii*. The figure was generated using *Loupe Browser*.

**Supplementary Figure 13.**
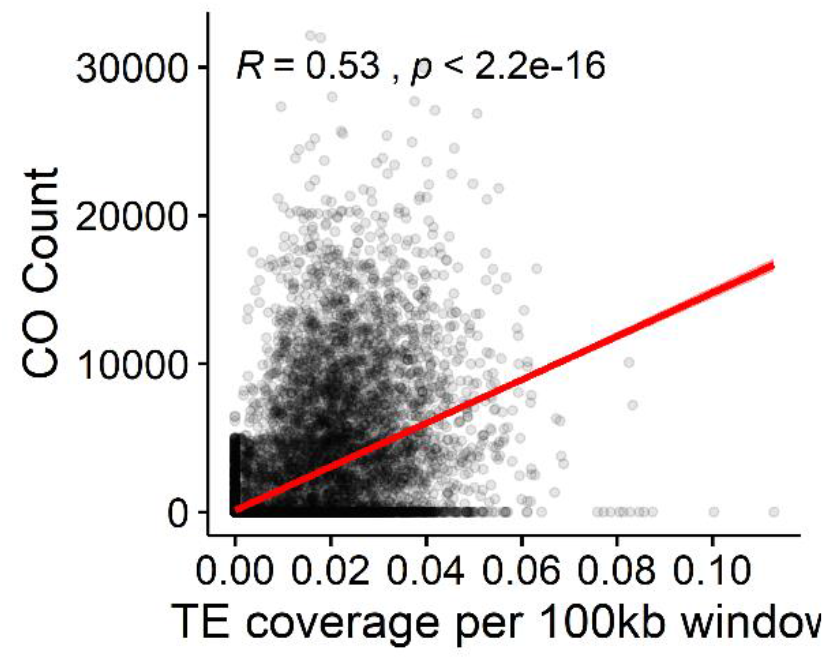
TE and CO correlation. Spearman’s rank correlation of crossover count and DNA transposons (*Stowaway* and *Tip100*) coverage in a sliding genome window. Each dot indicates a window.

**Supplementary Figure 14.**
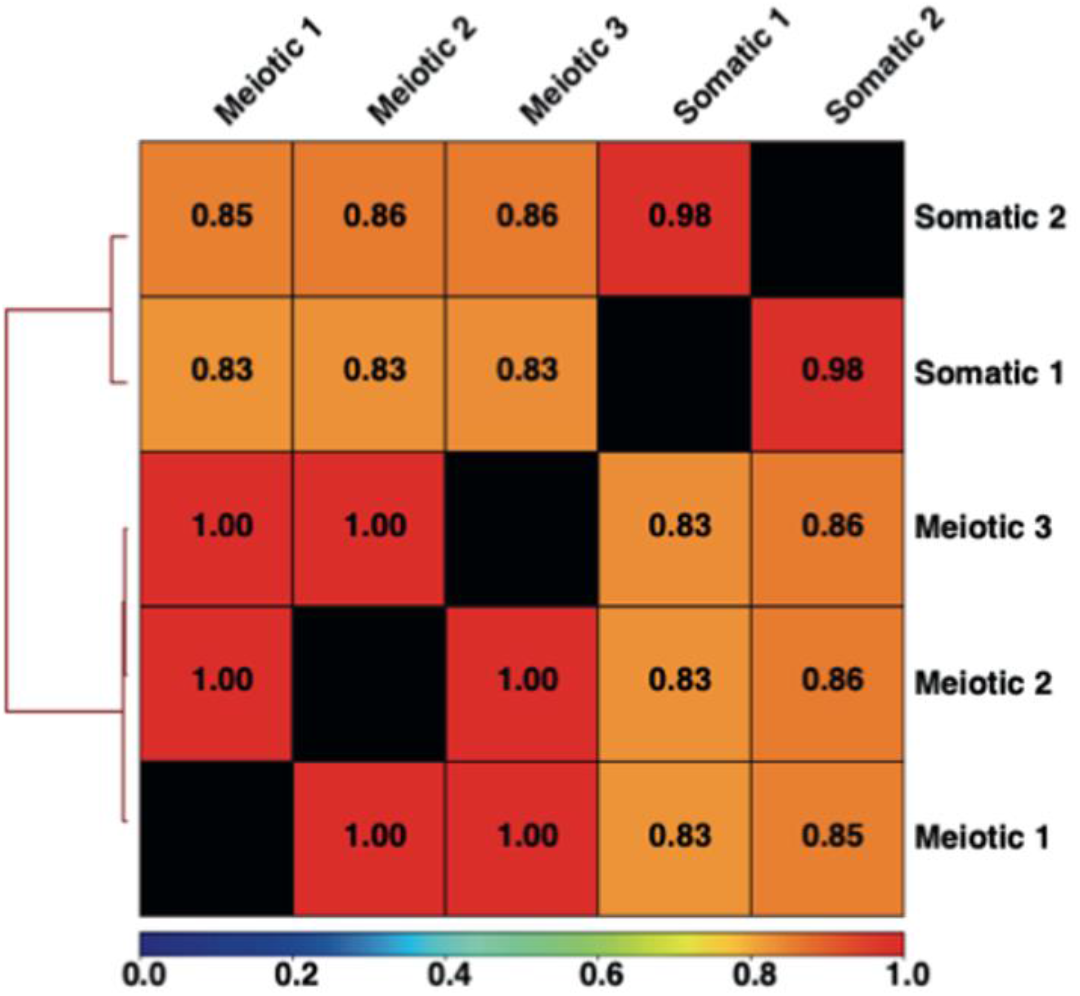
Pearson correlation of ACRs. Comparison of read distribution over the genome between tissues and between biological replicates.

**Supplementary Figure 15.**
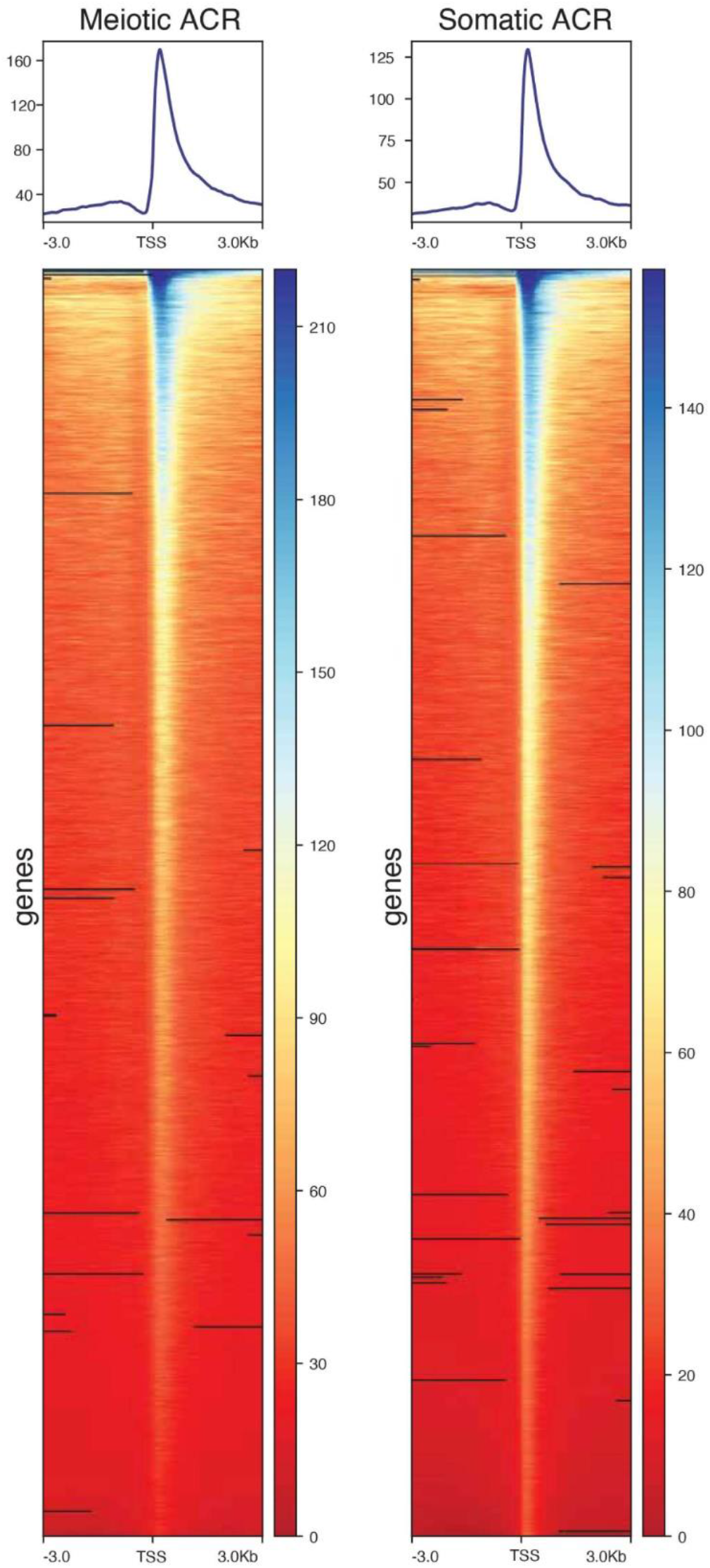
ATAC-seq peaks at transcription start sites (TSS). Read density across ATAC-seq peaks at TSS and their 3kb flanking regions. Each row of the heatmap represents one gene. Coverage is normalized in RPKM.

**Supplementary Figure 16.**
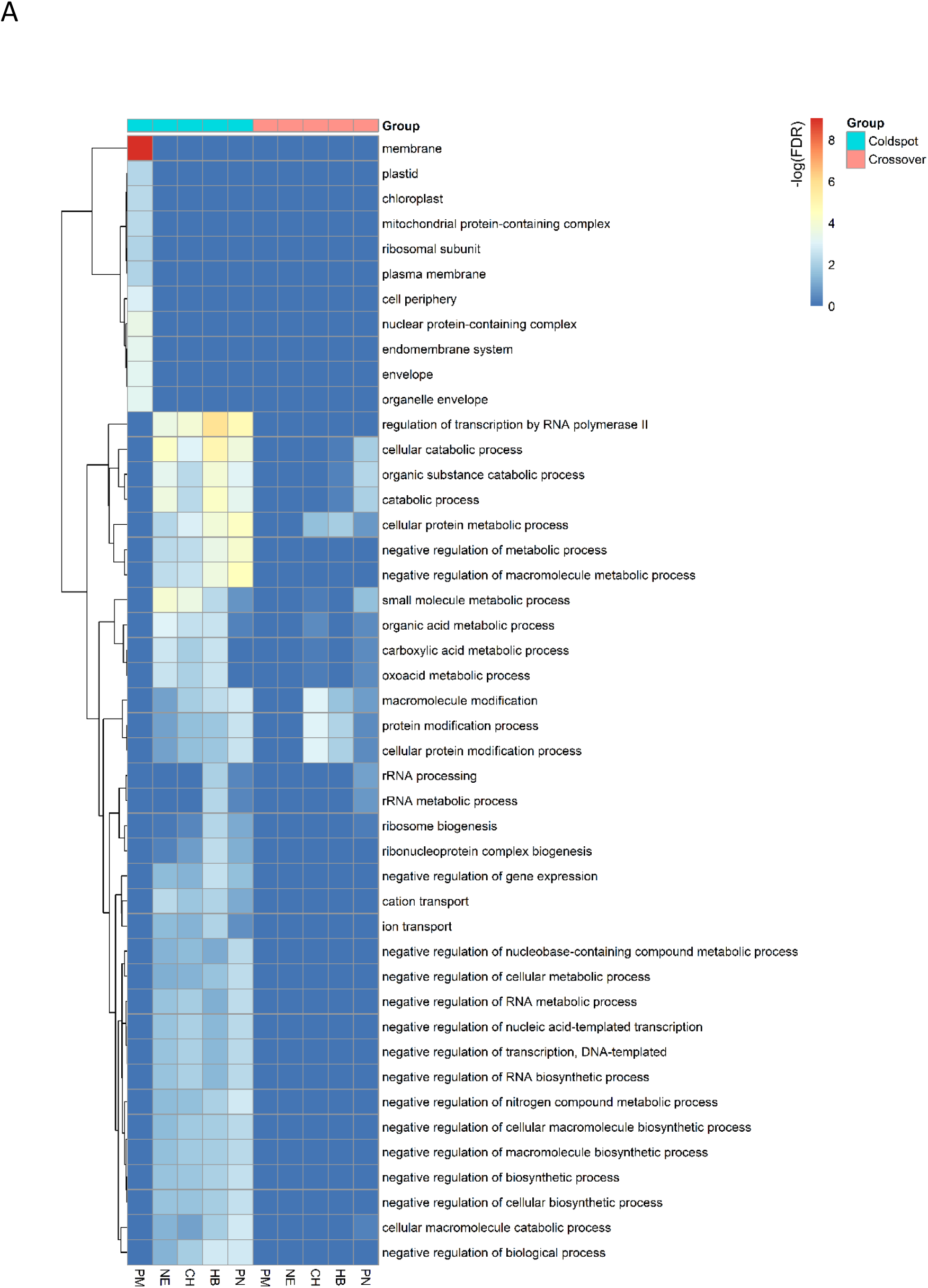
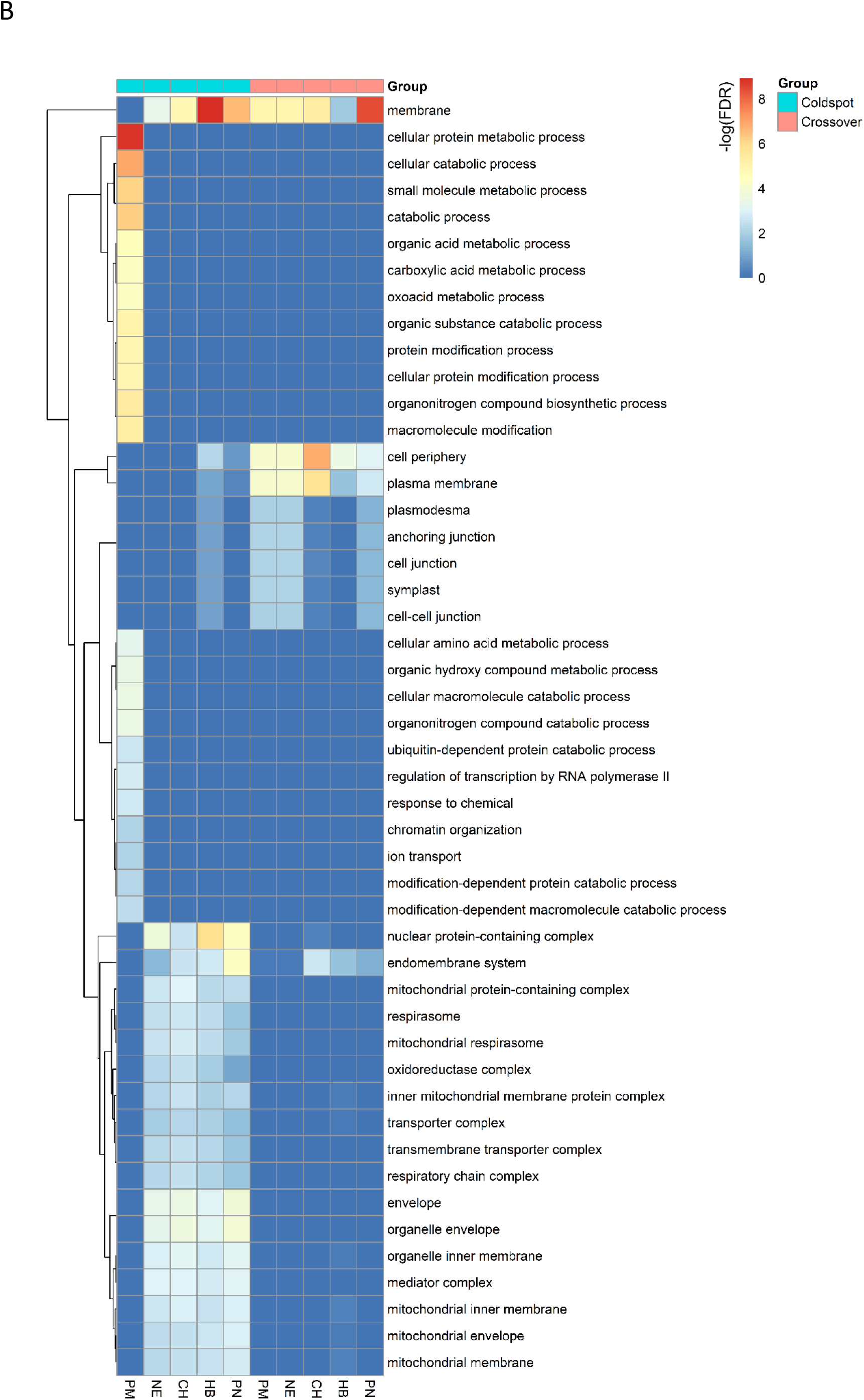
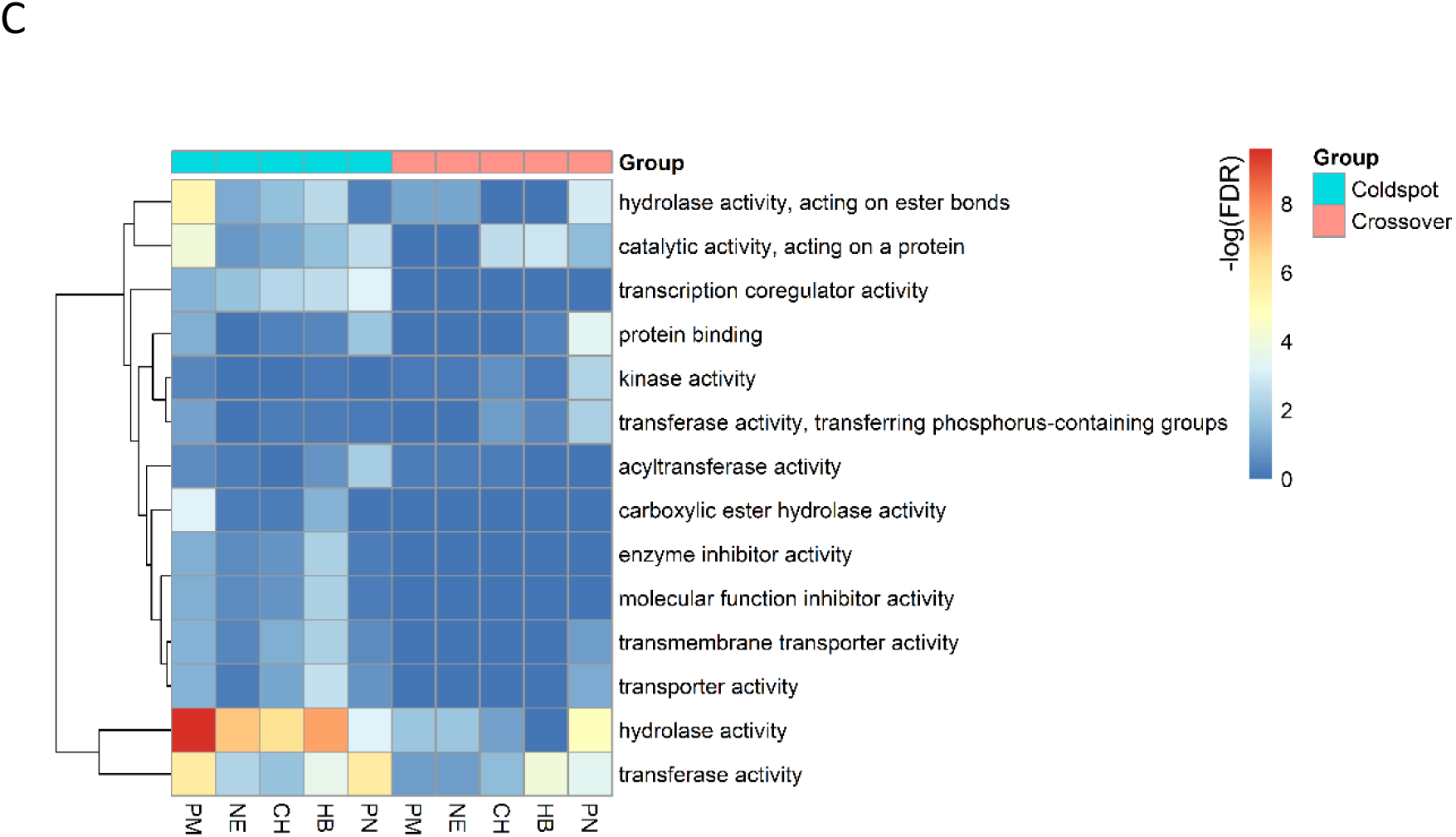
Functional enrichment in coldspot and CO regions. The minimum fold enrichment is 1.5. We separately reported the overrepresented terms by category: biological process (A), cellular location (B) and molecular function (C).

**Supplementary Figure 17.**
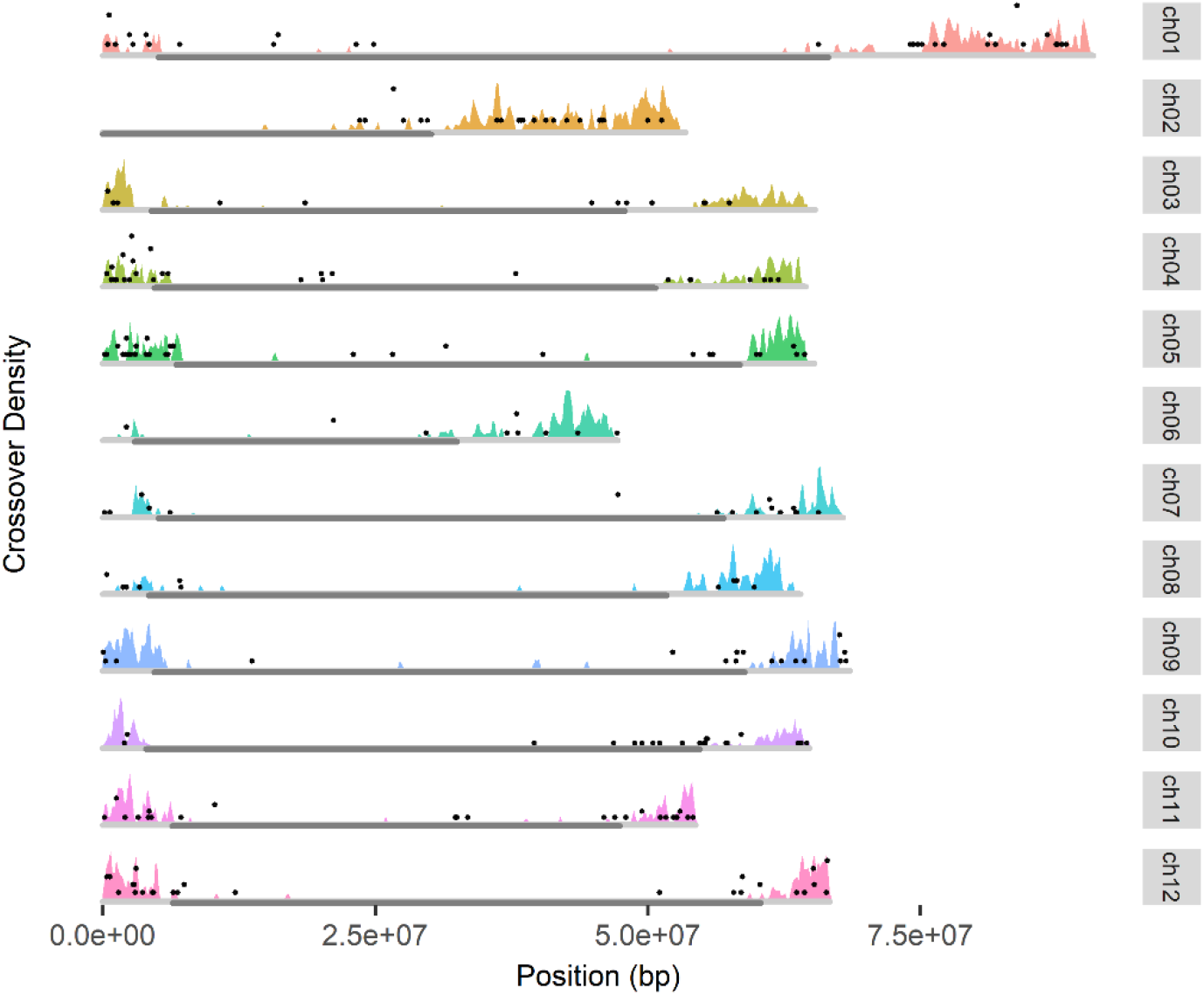
Resistance genes across the tomato genome. The black dots representing the frequency of R genes is plotted with the recombination landscape of the *S. lycopersicum* x *S. pennellii* hybrid

